# Humans use a dual policy to improve inferences during epistemic information seeking

**DOI:** 10.1101/2025.10.08.681186

**Authors:** Yinan Cao, Clémence Alméras, Junseok K. Lee, Inès Maye, Valentin Wyart

**Author notes:** Correspondence (Y.C.), (V.W.). Co-first author (Yinan Cao, Clémence Alméras).

## Abstract

Everyday decisions aim not only to earn rewards but also to learn about the world. Across three studies (total *N* = 702), we examined how people gather epistemic information stripped of rewarding value, and compared their strategy with reward seeking in otherwise matched conditions. Computational modeling of human behavior revealed a two-stage information-seeking policy, where participants first repeatedly sample each novel option in turn to test provisional hypotheses, a process we call ‘streaking’, before transitioning to uncertainty-guided exploration. While artificial neural networks trained to optimize inference accuracy acquired uncertainty-guided exploration but not early streaking, this two-stage policy improves human inference accuracy under noisy belief updating. Streaking and uncertainty-guided exploration tend to be co-expressed in the same individuals but map onto distinct psychological traits. Together, these results offer a novel account of human information seeking, clarifying its motives and benefits in epistemic contexts beyond the reward-centric explore–exploit tradeoff.

## Introduction

Many everyday choices pit the safety of the familiar against the lure of the unknown^1,2^. Should we buy our favorite snack from the vending machine or risk trying something new? Psychologists and neuroscientists capture this “explore–exploit dilemma”^3^ using bandit tasks, in which observers maximize total reward by sequentially sampling from options with different payoff distributions^4–6^. The sequence of choices is determined by the observer’s *policy*, that is, the systematic strategy that maps the current state of knowledge to an action. A policy may favor exploiting the best-known option or exploring to reduce uncertainty about alternatives. Typically, exploration is framed as a policy-level tradeoff with immediate reward, whereby short-term gains are sacrificed to learn about other options. Even with delayed outcomes, the information gathered still serves prospective valuation, i.e., computing expected future payoff during sampling, much like consulting restaurant reviews before choosing where to eat^7–11^.

But information in and of itself can motivate behavior^12^. Influential laboratory-based work shows that rodents and primates seek cues that reduce uncertainty about forthcoming rewards even when those cues do not alter reward contingencies^13–17^. Yet the information there still concerns the reward structure of the environment, reducing uncertainty about when or how much reward will arrive. Moreover, the informative choice made in controlled experiments is often single-shot rather than sequential, which limits what we can learn about policies that guide multi-step information gathering. Besides, many everyday forms of real-life exploration are epistemic (knowledge acquisition), with policies unfolding over multiple steps. Their goal is to build internal models of the environment independently of immediate, or even prospective valuation. Children’s play, learning the syntax of a new language, or orienting oneself in an unfamiliar city all involve sampling information that is not tied to payoff, yet provides structural knowledge that supports flexible behavior^18–20^. How releasing sampling from prospective valuation shapes information-seeking policies is much less known.

Here we use a sequential sampling task that dissociates reward-driven from epistemic exploration and test it in a large sample of participants. In the epistemic sampling condition, participants showed an overall stronger tendency to sample more uncertain options (uncertainty-guided sampling), which occurred after a period of *early streaking*, i.e., repeatedly sampling the same novel option in short streaks before sampling another one. Although statistically suboptimal, streaking co-occurred in participants with more effective uncertainty-guided sampling and with higher overall decision accuracy. We built a computational model that captures the full repertoire of observed policies and formalizes streaking as a *local*, threshold-based cognitive phenotype alongside *global* uncertainty-guided sampling. Importantly, model simulations showed that both streaking and uncertainty-guided sampling provide epistemic benefits in the presence of noisy belief updates—characteristic of human cognition.

Investigating individual differences in participants’ policy revealed a dissociation between the cognitive traits that relate to early streaking and uncertainty-guided sampling. Need for cognitive closure predicted streaking tendency, whereas general reasoning ability predicted effective uncertainty-guided sampling. Artificial neural networks trained to optimize epistemic accuracy acquired near-optimal uncertainty-guided sampling but failed to acquire early streaking. This distinction underscores early streaking as a distinctively human strategy, shaped by human cognitive constraints not shared by artificial neural networks. This pattern of findings delineates a schema in which normative global policies are complemented by local sampling rules shaped by traits and the limited precision of neural computations.

## Results

### Task design

Adult volunteers (see Methods) performed a sequential sampling task (Fig. 1). They sampled from two options depicted as bags, each containing colored gems of a certain shape (e.g., triangle-shaped gems for one bag, pentagon-shaped gems for another bag). Each bag contained gems ranging from blue to orange, although one color predominated. For example, the bag of pentagonal gems could contain mostly orange ones. In each game of the task, consisting of 8–20 sampling trials, participants were presented with two novel bags, whose dominant colors were set independently from each other (so that sampling from one bag provided no information regarding the dominant color of gems in the other bag). Participants were asked to sample gems from the two bags, one gem at a time. On each trial, picking a bag (by choosing its gem icon) revealed the color shade of a gem drawn from that bag (Fig. 1A).

**Figure 1.**
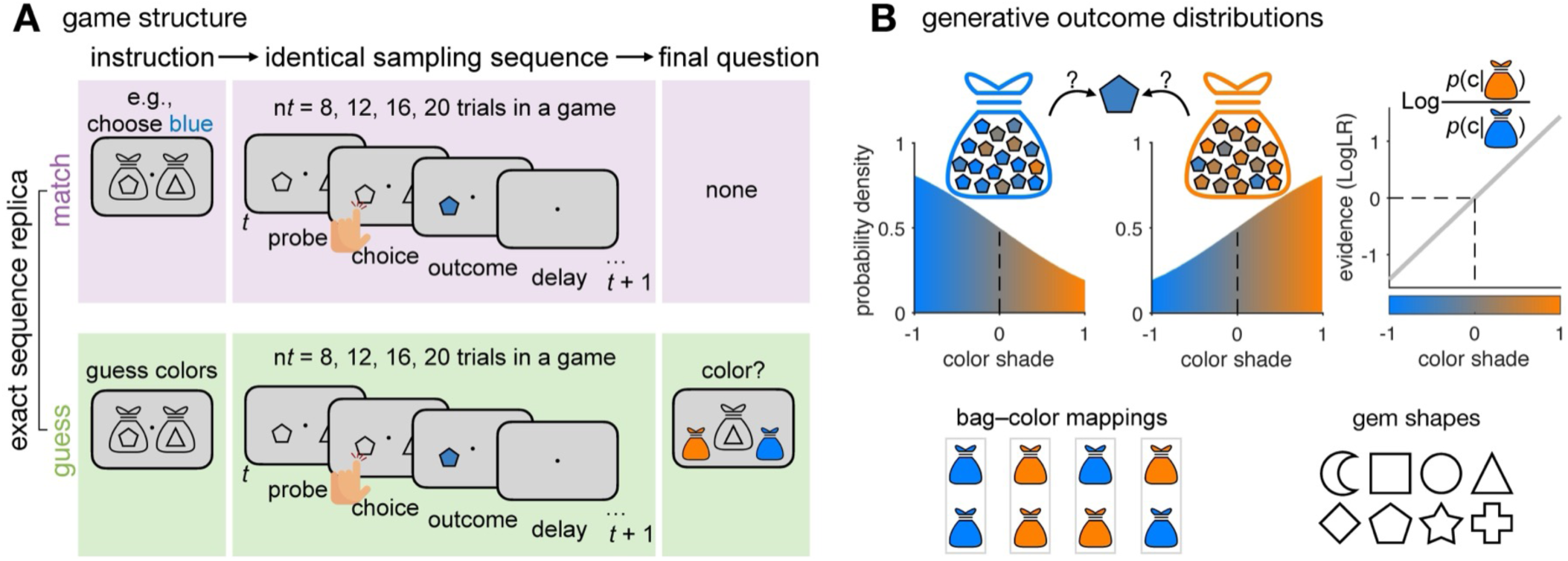
Experimental paradigm. (**A**) In each game, participants chose between two options (bags), each *independently* associated with either predominantly blue or orange gems (depicted as different shapes). The dominant color of each bag was not directly observable and had to be inferred through trial-and-error sampling. In the MATCH condition, participants learned to select the gems matching a target color (for example, blue). In the GUESS condition, participants were asked to sample the options to learn the mapping from gem shape to its bag’s dominant color. Both conditions included a sampling phase of 8, 12, 16, or 20 choices (one choice per trial). The length of this phase varied randomly across games and was not disclosed before each game. In the GUESS condition only, participants reported after the sampling phase the dominant color associated with one of the two bags, selected at random. (**B**) Outcomes (color shades) were drawn from generative distributions (Methods). For the blue bag (the bag with predominantly blue gems), approximately two thirds of outcomes were blue and one third orange (error rate ∼1/3), with the reverse for the orange bag. Each outcome conveyed a log-evidence (log-likelihood ratio, logLR) favoring the orange- vs. blue-bag hypothesis; cumulative logLR across trials gave the posterior belief (Methods). Associations between bags and dominant colors were independently assigned so that both options could map to the same or different dominant color(s).

In one condition (MATCH), participants were instructed to sample gems matching a target color (e.g., blue). The goal was to identify and repeatedly select the option (bag) most consistent with the target dominant color, thereby maximizing the number of target-colored gems collected. This instruction thus assigned rewarding value to colors and required participants to trade off exploration against immediate reward maximization. As in standard ‘bandit’ tasks^4,11^, participants’ performance was evaluated by the match between sampled gems and the target color throughout the game. In another condition (GUESS), there was no target color. Participants were asked to sample the options to learn the dominant gem color associated with each bag. Each game ended unpredictably after 8–20 samples. Participants then indicated the dominant color associated with one of the two bags, selected at random. In this condition, participants’ performance was evaluated solely by the accuracy of this final decision regarding the dominant color of the probed option. This GUESS condition therefore divorced exploration from any form of immediate or prospective reward maximization.

Apart from the instructions, the two conditions were identical, allowing direct comparison of sampling patterns. Of note, a single outcome (color shade) could not fully reveal the dominant gem color of the associated bag, since outcomes were generated from probability distributions (Fig. 1B). Participants could not infer the dominant color of one bag from the dominant color of the other bag, because bag–color associations were independent across options. After predefined data quality control, 420 participants were included in the analyses (see Methods). We analyzed participants’ choice patterns to identify their information-seeking strategies (policies) in the two conditions. We begin with model-free comparisons of behavioral patterns across conditions, then use computational approaches—fitting cognitive models to human data and training recurrent neural networks with different objectives—to identify the latent cognitive processes underlying the observed patterns, and finally examine individual differences in human information-seeking policies and their relation to performance.

### Behavioral performance and information-seeking policy

We first assessed participants’ performance in each condition (Fig. 2A). As expected, in the MATCH condition, participants reliably selected the target-colored option by the last trial of each sequence (mean last-trial accuracy 0.786, 95% CI [0.770, 0.802]), significantly above chance, t(419) = 34.52, p < 0.0001. Performance further improved with sequence length, t(419) = 2.33, p = 0.021, with an average slope of 0.0025 [0.0004, 0.0045]. In the GUESS condition, participants correctly identified the option–color mapping above chance at the final probe (mean accuracy: 0.785 [0.774, 0.797]), t(419) = 48.96, p < 0.0001, and accuracy showed a clearer increase with sequence length, t(419) = 12.34, p < 0.0001, mean slope 0.0104 [0.0088, 0.0121] (Fig. 2A).

**Figure 2.**
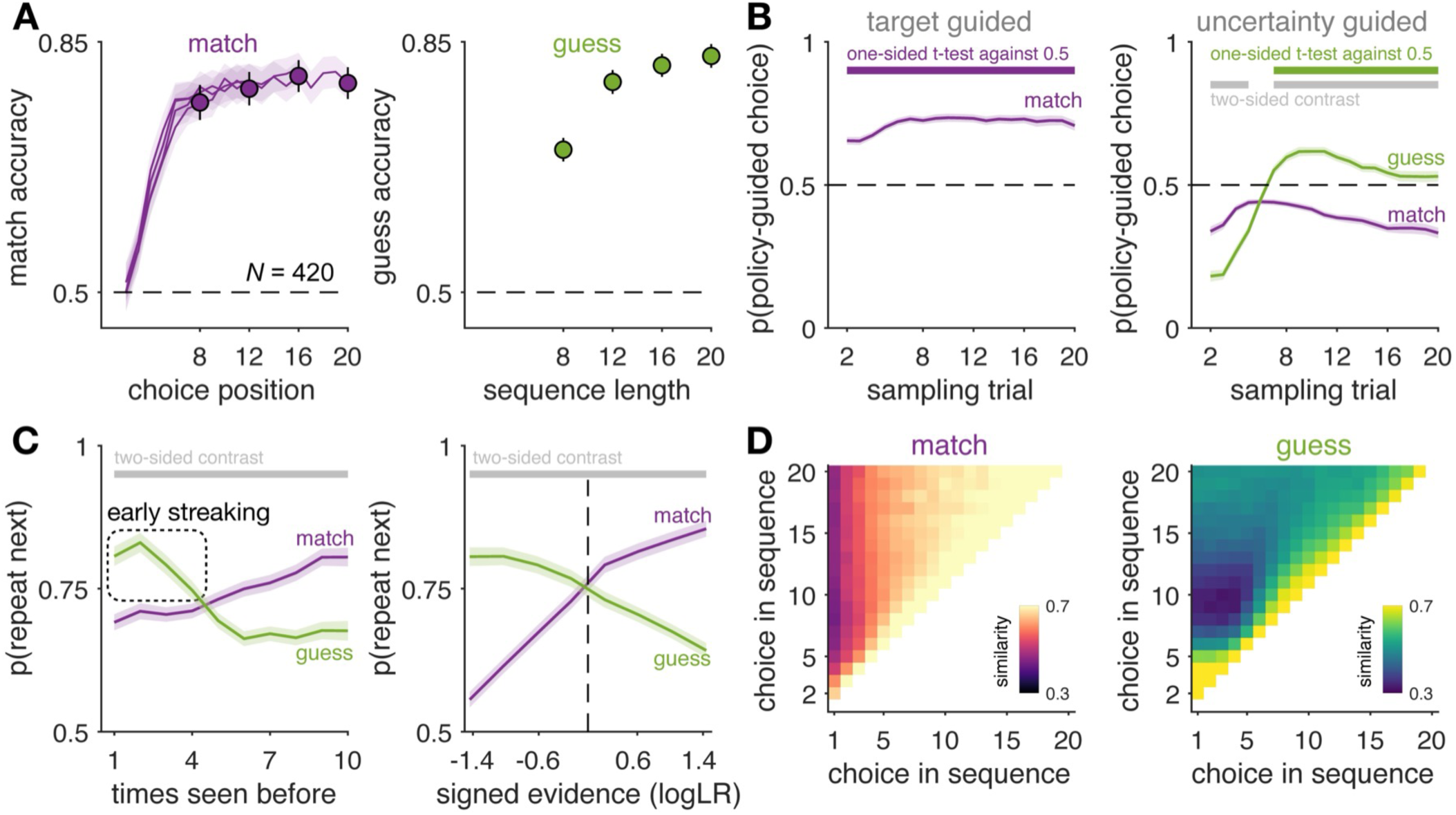
Human behavior. (**A**) Accuracy in the two conditions. In the MATCH condition, accuracy is the probability of selecting the option associated with the target color in games with counterfactual hidden structure (one bag linked to orange, the other to blue). In the GUESS condition, accuracy is the correctness of the final probe after each game. (**B**) Fraction of choices aligned with directed-exploration policies across sampling trials. On each trial, based on the outcomes observed so far, we identified the most target-like option as the one with the highest accumulated evidence toward the target color, and the most uncertain option as the one with the lowest absolute accumulated evidence. Thick lines indicate statistically significant effects (one-sided t-test for policy alignment, FDR < 0.005) and between-condition contrasts (two-sided t-test, FDR < 0.005). (**C**) Left: probability of repeating a choice as a function of how many times the current option has been sampled. Right: probability of repeating a choice as a function of signed evidence, expressed in logLR unit. In the MATCH condition, signed evidence reflects target-aligned evidence; in the GUESS condition, it reflects belief-aligned evidence, assuming an accumulation of trial-by-trial evidence for each option given the actual choices (Methods). Error bars show 95% confidence intervals (CIs) across 420 participants. (**D**) Mean choice-to-choice similarity within the sampling phase, pooled across sequences of lengths 8, 12, 16, and 20.

We next examined whether participants’ sampling behavior was driven by expected reward and uncertainty. On each trial based on the running sum of color evidence, we classified the objectively most target-like option (in the MATCH condition, the option with highest accumulated evidence toward the target color) and the most uncertain option (in both conditions, the option with lowest absolute accumulated evidence). As expected from the task structure, choices in the MATCH condition predominantly favored the target-like option (one-sample t-test against 0.5 across trials, peak t(419) = 35.19, p < 0.0001), indicating reward-guided sampling. By contrast, choices in the GUESS condition showed a significantly stronger bias toward the most uncertain option (paired t-test between conditions, trials 7−20, FDR-corrected p < 0.005; peak t(419) = 20.51, p < 0.0001). These patterns reveal systematic sampling policies wherein participants adopted reward-guided sampling in the MATCH condition and uncertainty-guided sampling in the GUESS condition (Fig. 2B). Of note, the probability of policy-guided choices remained below 1 (Fig. 2B), indicating imperfect policy alignment.

Closer inspection also revealed a systematic deviation from the uncertainty-guided policy early in GUESS games (around trials 2–5), where participants showed a temporary bias toward the relatively more certain option. What explains this early behavior? Visual inspection of choice sequences suggested that participants were repeatedly sampling the same option in short “streaks” before switching, for example, choosing AAAAABBBBB rather than alternating. We term this pattern *streaking*, an initial local form of information seeking in which participants concentrate early samples on one option at a time.

To characterize this early streaking policy, we examined how the probability of repeating a choice depended on how many times that option had already been sampled. In the GUESS condition, participants showed a stronger tendency to resample options they had sampled only once or twice (t(419) > 17.17, p < 0.0001; Fig. 2C, left), revealing a strong repetition bias early in the game. Choice-to-choice similarity matrices confirmed this pattern (Fig. 2D): participants showed high choice similarity among early trials (e.g., trials 1–5), followed by a shift toward the other option (e.g., trials 6–10), resulting in low similarity between these two streaks—a pattern resembling AAAAABBBBB choice sequences. By contrast, this streaking pattern was absent in the MATCH condition, where participants progressively converged on the most rewarding option. Separately, both conditions exhibited a general repetition bias throughout the sampling phase, visible as elevated similarity between adjacent trials across the entire choice-to-choice similarity matrix (bright band just above the diagonal; Fig. 2D).

Complementary analyses further clarified the decision dynamics driving these sampling behaviors. In the GUESS condition, participants were more likely to resample an option when the current outcome conflicted with their emerging belief about its hidden color, whereas in the MATCH condition, resampling was more likely when the outcome aligned with the target color, consistent with reward-maximizing behavior (Fig. 2C, right). Time-locked analyses of the first switch within each game provided stronger evidence for this pattern (Supplementary Fig. S1). In the GUESS condition, participants made their first switch only after receiving a large belief-confirmatory outcome (evidence supporting their current hypothesis about that option). By contrast, in the MATCH condition, participants switched after a large target-conflicting outcome (evidence that the current option was not rewarding). Together, these results support an evolving hypothesis-checking strategy in the GUESS condition, where participants initially test one option via streaking and switch only once sufficient confirmatory evidence has accrued for that option.

In sum, despite identical generative statistics, the GUESS and MATCH conditions elicited distinct choice policies. In the MATCH condition, choices favored the option most likely to be aligned with the target color, whereas in the GUESS condition, choices favored the option whose hidden dominant color was most uncertain. In the GUESS condition, however, this choice policy emerged only after a phase of early streaking, corresponding to initial bouts of repeated choices from each option in turn.

We further corroborated these behavioral patterns in two independent experiments (*N* = 282 participants in total), demonstrating replicability with distinct stimuli (Supplementary Fig. S2) and generalizability to a ternary-choice setting (Supplementary Fig. S3). In the ternary-choice study, sampling patterns indicate that participants in the GUESS condition engaged in the same two-stage policy, initially streaking through each of the three options *in turn* before transitioning to uncertainty-guided exploration (see Supplementary Fig. S3D).

### Computational model of sequential sampling

To formalize the observed sampling policies, we developed a computational model of sequential sampling in the MATCH and GUESS conditions, building on and extending the foundational model from our previous work^21^. We first define an ideal observer to serve as a normative reference, and then describe how our model extends beyond this reference to reflect human sampling by including a threshold rule and imprecise belief updating.

#### Normative benchmark

A Bayes-optimal agent should accumulate evidence trial by trial as log-likelihood ratios (logLR) for each option. In the MATCH condition, it selects the option most aligned with the target color (most target-like), which maximizes expected reward. In the GUESS condition, where no target is defined, it selects the option with the smallest absolute accumulated evidence (most uncertain), which maximizes the expected accuracy of inference when the game can terminate at any moment. These are the optimal sampling policies of an ideal observer.

#### Suboptimal evidence accumulation model

While the ideal observer provides a critical normative benchmark, it does not account for the idiosyncrasies of human sampling behavior. Our extended suboptimal model flexibly instantiates the normative policies with two sensitivity parameters (Fig. 3A): *β*_c_ for uncertainty-guided sampling and *β*_v_ for reward-guided sampling, with *β*_v_ not defined in the GUESS condition (because there is no target in GUESS). In the limit 𝛽 → +∞ the policies are optimal, and free *β* parameters make the model flexible (see Supplementary Fig. S4 for illustrative model simulations).

**Figure 3.**
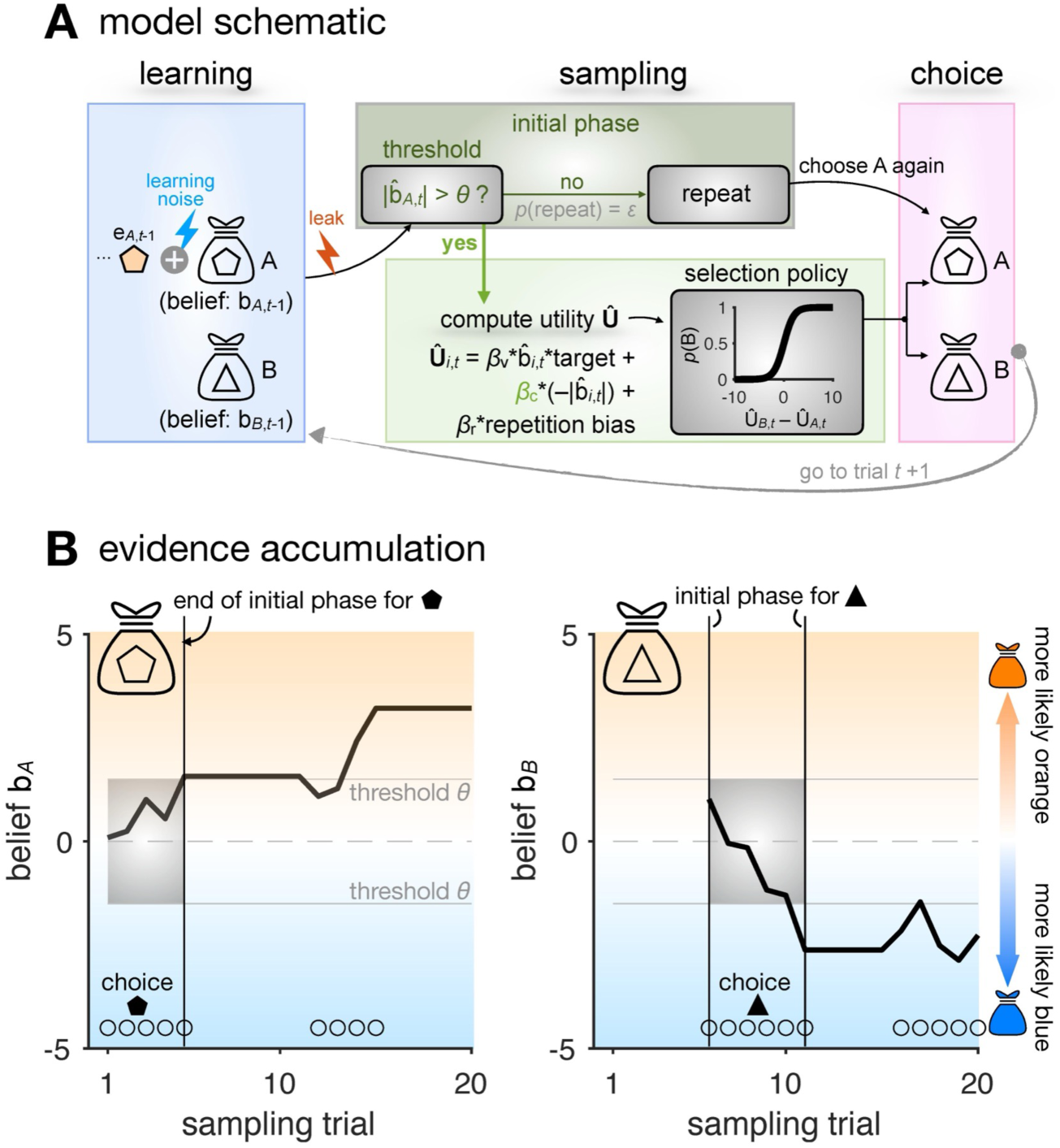
Computational model. (**A**) Learning was modeled as a noisy and leaky integration of evidence over trials. Choices followed a softmax over a utility that combined three components: (i) target-aligned accumulated evidence (relevant in the MATCH condition only), weighted by a target-value sensitivity parameter (𝛽ᵥ); (ii) belief uncertainty, weighted by an uncertainty sensitivity parameter (𝛽c); and (iii) a general repetition bias, weighted by a repetition sensitivity parameter (𝛽r). An initial sampling phase preceded the utility-based sampling stage. During this initial phase, choices repeated (with probability *ε*) without relying on the utility until the accumulated evidence for the current option crossed a threshold *θ*. (**B**) For an example sequence, cumulative evidence for each option equals the evolving posterior belief about the bag’s dominant color. The initial sampling phase ends when the belief crosses the threshold *θ*.

In addition, we incorporated a local, threshold-based streaking mechanism controlled by a confidence level *θ* and a repetition strength *ε*. This mechanism drives an initial sampling phase where the agent repeats the same choice until the accumulated evidence for that option exceeds the threshold *θ*. Together, *θ* and *ε* determine the agent’s probability of resampling the same option on successive trials (Fig. 3 and Supplementary Fig. S4).

Finally, the model included a general repetition bias parameter (*β*_r_) for choice autocorrelation^22^ beyond the early streaking phase, a leak parameter (*δ*) for gradual information loss across trials, and a learning-noise term (*σ*) for imprecise belief updates. These extensions capture the leaky and noisy nature of human belief updating^23,24^. Parameter and model recovery confirmed that the full model and its components were identifiable^25,26^ (Supplementary Figs. S5–S6).

### Model fits and parameter estimates

We fitted the model to each participant’s choices (see Methods)^27^ to assess how well it captured human behavior. Model simulations with the best-fitting parameters reproduced the human sampling patterns (Fig. 4A). These included reward-guided sampling in the MATCH condition, stronger uncertainty-guided sampling in the GUESS condition, and early streaks of repeated choices in GUESS but not in MATCH. The model also reproduced the divergent resampling tendencies as a function of signed evidence across conditions: in the MATCH condition, participants were more likely to resample the same option when evidence was consistent with the target color, whereas in the GUESS condition resampling increased under belief-violating evidence. Together, the generative performance of these simulations confirms that the model accounts for both the global and local policies of human sampling.

**Figure 4.**
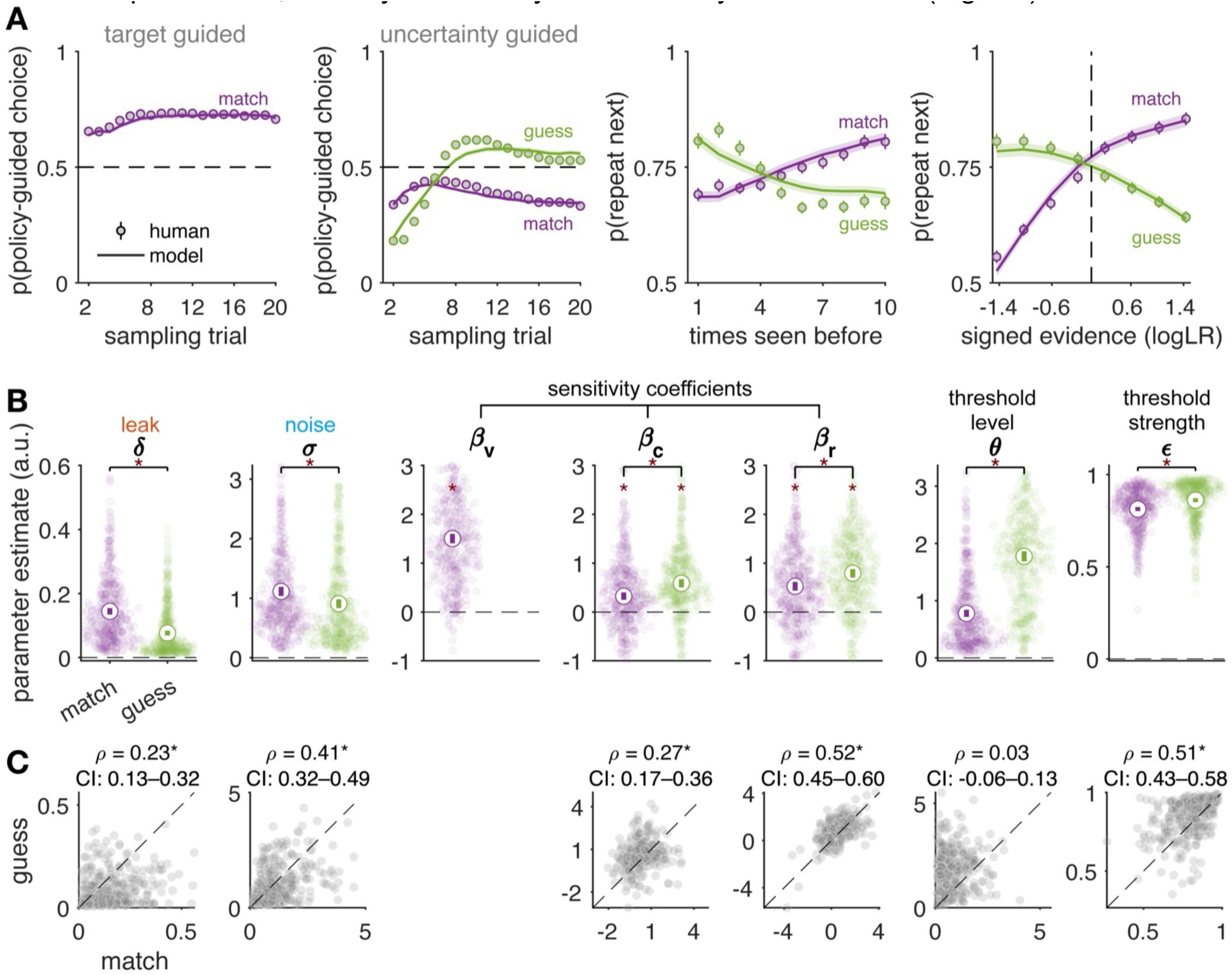
Model fits and parameters. (**A**) Model simulations based on the suboptimal evidence accumulation model using best-fitting parameters. (**B**) Estimated best-fitting model parameters. Asterisks indicate statistically significant contrasts between the MATCH and GUESS conditions (paired-sample t-tests, two-sided), as well as significant deviations from zero (one-sample t-tests within conditions, two-sided), both assessed using max-statistic permutation tests with family-wise error correction (FWE < 0.05). (**C**) Parameter correlation across conditions. Asterisks indicate significant Spearman correlations between the MATCH and GUESS conditions, assessed via bootstrapping (5,000 resamples with replacement). Bootstrap means and 95% CIs are shown for each correlation estimate.

All sensitivity parameters were significantly above zero in their respective conditions (t(419) ≥ 8.51, p values < 0.001), and they differed between conditions where defined (|t(419)| ≥ 5.01, FWE-corrected p values < 0.001). In particular, uncertainty sensitivity (*β*_c_) was markedly higher in the GUESS than in the MATCH condition (t(419) = 5.14, FWE-corrected p < 0.001), as were the threshold parameters *θ* and *ε* (t(419) > 8.64, FWE-corrected p values ≤ 0.0002; Fig. 4B), consistent with stronger early streaking and stronger uncertainty-guided sampling in GUESS games. These effects were replicated in an independent ternary-choice study (*N* = 243; Supplementary Fig. S3E). Together, these findings corroborate that the global policies governed by sensitivity parameters, and the local streaking policy governed by threshold parameters, were systematically modulated by task condition (Fig. 4B).

In addition, learning parameters were also affected by conditions. Learning noise *σ* and leak *δ* were significantly lower in GUESS games than in MATCH games. This difference suggests that shifting the task framing from reward seeking (MATCH) to information seeking (GUESS) was associated with: 1) reduced internal noise in belief updating, and 2) reduced working memory leak of accumulated evidence. Thus, instructions modulated the fidelity of belief updating in addition to shaping global and local sampling policies. Across participants, *β*_r_ correlated positively between the MATCH and GUESS conditions (bootstrap mean *ρ* = 0.52; Fig. 4C), indicating a general switch cost common to both conditions.

### Contributions of leak and noise to suboptimal decisions

Beyond capturing global and local sampling policies, our model fits also revealed systematic differences in learning noise and leak between conditions. We therefore asked whether these imperfections quantitatively accounted for deviations from optimality. Bayesian modelcomparison showed that the models including learning noise explained participants’ sampling behavior decisively better than those without the noise term (Fig. 5A, B and D, E; family-wise comparison, noise vs. no noise, exceedance probability P_exc_ > 0.99). This held in both conditions. In addition, the MATCH condition featured a strong contribution of memory leak: the full model including both leak and noise best described sampling in this condition (P_exc_ > 0.99). By contrast, in the GUESS condition, learning noise emerged as the dominant source of suboptimality (Fig. 5D; Supplementary Fig. S7). Of note, randomness from the softmax rule, governed by the sensitivity parameters *β*, also contributed to the overall variability in sampling choice. Learning noise captured an additional, independent source of variability beyond this selection stochasticity (see parameter recovery in Supplementary Fig. S6).

**Figure 5.**
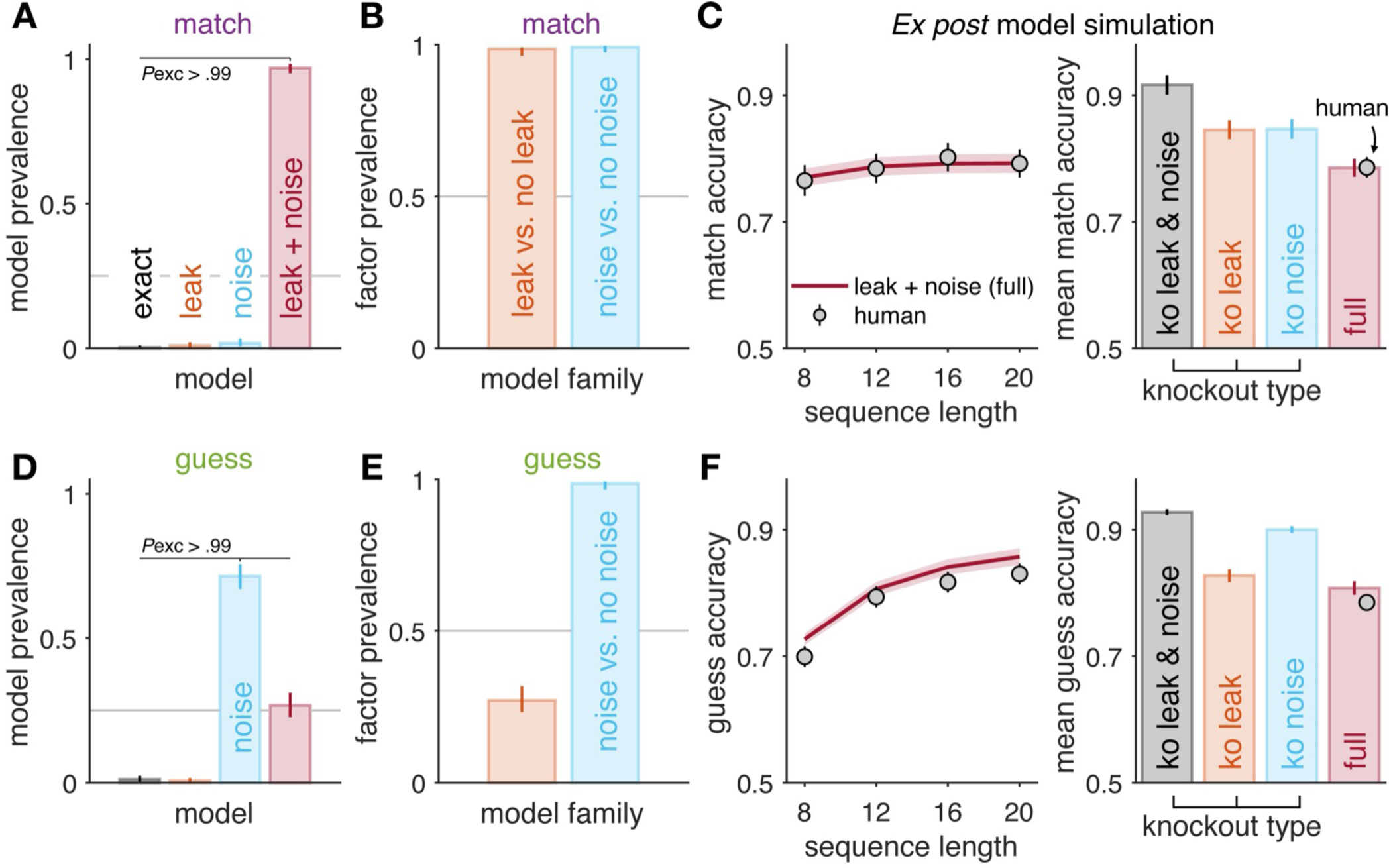
Model comparison and simulations. Top row: MATCH condition (A–C). Bottom row: GUESS condition (D–F). (**A**, **D**) Posterior model frequencies from random-effects Bayesian model selection. Models: exact (no memory leak and no learning noise; other parameters retained, including sensitivity coefficients and threshold parameters), leak only, noise only, and leak + noise (full). (**B**, **E**) Family-wise comparisons testing the independent contribution of leak and learning noise to models’ goodness-of-fits. (**C**, **F**) Left: model-predicted accuracy vs. human data. Right: Effect on model-predicted performance of removing (“knocking out,” i.e., “ko”) individual factors (leak, noise, or both). Accuracy in MATCH condition (top row): correctly choosing the target-aligned option; Out-of-sample inference accuracy in GUESS condition (bottom): correctly judging dominant gem color after sampling phase. Model parameters were estimated from sampling behavior only; predictions of guess accuracy were therefore “out-of-sample” with respect to the fitted domain. Error bars indicate 95% CIs.

To assess how leak and noise each affected behavior, we performed “knockout” simulations, setting each parameter to zero while keeping others at fitted values (Fig. 5C, F). In the MATCH condition, both knockouts produced similar deviations from the full model (Fig. 5C), indicating equal contributions to suboptimality. In the GUESS condition, however, only removing noise, not leak, caused substantial deviation (Fig. 5F, right), suggesting that noise primarily drove inference suboptimality.

In sum, participants’ sampling reflected a dual, two-stage strategy combining near-optimal global policies with a local, structured strategy. At the global level, sampling was guided by reward in the MATCH condition and uncertainty in the GUESS condition, approximating normative policies but constrained by the condition-specific computational limits. At the local level, an early streaking phase appeared specifically in the GUESS condition.

### Recurrent neural networks acquire uncertainty-guided sampling but not early streaking

Human behavior showed a dual architecture, combining near-optimal global sampling policies with a local streaking strategy. This raises a central question: Do these policies emerge naturally from optimization objectives alone? To investigate this, we trained standard recurrent neural networks (RNNs)—with objectives specifically matched to each task condition—on the same task performed by human participants.

In the MATCH condition, RNNs trained via a reward-maximizing policy-gradient method converged on strong reward-guided sampling, closely paralleling human behavior (Fig. 6C). Principal component analysis of hidden activity showed that the network organized its representations primarily along a low-dimensional axis (Fig. 6D), with effective dimensionality around 1.15.

**Figure 6.**
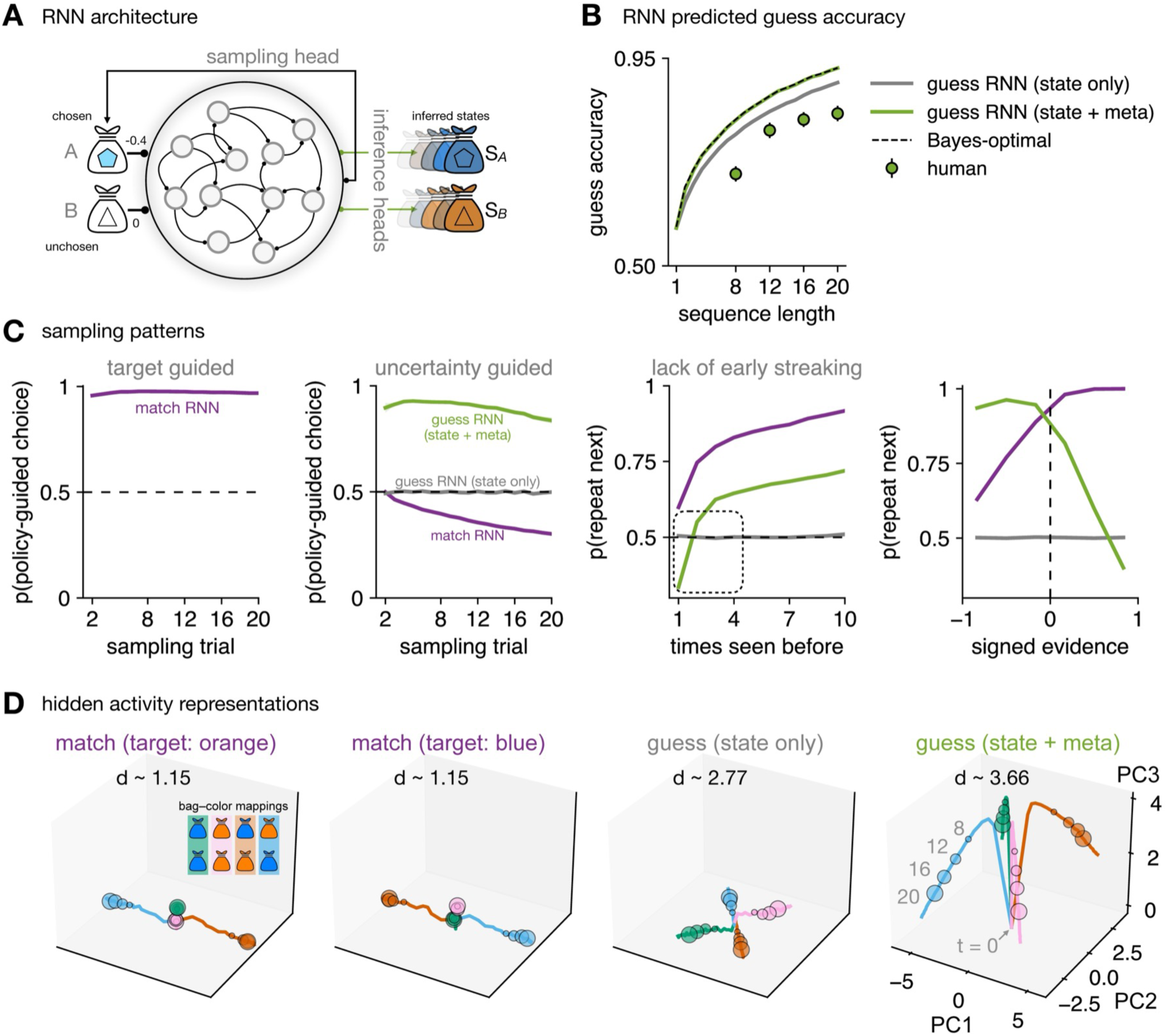
RNNs, sampling patterns, and representational geometry. (**A**) The RNN receives input conditioned on the previous sampling choice (e.g., if option A is chosen, the input is the color shade corresponding to A’s outcome, while the unchosen option B yields an ambiguous zero outcome), combined with the previous recurrent activity. Outputs include one sampling head for action selection and two inference heads for state prediction. (**B**) GUESS accuracy was computed by simulating each game, taking the RNN output on the final trial after the last update to the chosen option, converting it to a categorical response (orange if > 0, blue if < 0), and scoring a match to the true target. Bayes-optimal: performance of an agent with perfect uncertainty-guided sampling (𝛽_c_→ +∞). (**C**) RNN-predicted sampling behavior. The MATCH RNN exhibits strong reward-guided sampling policy. Only the GUESS RNN with the meta-policy displays uncertainty-guided sampling; the state-prediction-only RNN does not. None of the RNNs reproduce the early streaking observed in humans. (**D**) Hidden activities of the MATCH RNN collapse onto a low-dimensional axis. By contrast, GUESS RNNs form separable representations of the four different bag–color mappings, and the addition of the meta-policy further enhances this separation, yielding a well-pronounced representational geometry. Curves: trajectories over sampling trials; circles: sequence lengths = 8 to 20 trials; ‘d’: effective dimensionality (Methods).

In the GUESS condition, we used the same RNN architecture but varied the training objective to predict hidden states of the games (bag colors; Fig. 6A). This network achieved high state-prediction accuracy (Fig. 6B) and formed well-separated representational geometries across different bag–color mappings (Fig. 6D, effective dimensionality 2.77; Supplementary Fig. S8A), yet converged on random sampling (*β*_c_ = 0; Fig. 6C; Supplementary Fig. S11). Even with extended training (Supplementary Fig. S9) and adequate hidden-layer capacity^28^, the network did not spontaneously discover uncertainty-directed exploration (Fig. 6C; Supplementary Fig. S10B). Maximizing predictive accuracy alone was therefore insufficient for the emergence of optimal sampling policies.

When trained with a hybrid objective—predicting hidden states and maximizing meta-cognitive certainty during sampling—the GUESS network achieved near-Bayes-optimal inference accuracy (Fig. 6B) and produced human-like uncertainty-guided exploration (Fig. 6C). The meta-policy objective, implemented as an auxiliary policy gradient loss, encouraged actions that reduced representational uncertainty. This meta-policy GUESS RNN showed higher effective dimensionality in the hidden layer (mean 3.662, min 3.503, max 3.795), with the third principal component encoding a moment-to-moment certainty signal (Supplementary Fig. S8B).

However, a critical discrepancy remained. None of the RNNs, from 4 to 64 hidden units, reproduced the early streaking policy observed in humans (Fig. 6C: 3^rd^ panel; Supplementary Fig. S10). Intriguingly, forcing streaked sampling at test time in the state-prediction RNN increased expected GUESS accuracy in late trials (Supplementary Fig. S11), suggesting benefits of streaking. However, the meta-policy RNN, having learned the uncertainty-guided optimum (𝛽_!_ → +∞), showed no marginal benefit from early streaking—even in late trials, as streaking would delay uncertainty-guided sampling—which is the Bayes-optimal policy in this condition. This likely reflects that once the RNN converges to the uncertainty-guided optimum, learning the streaking policy becomes trivial and thus not reinforced. These results suggest that streaking competes with optimal uncertainty-guided sampling in the meta-policy RNN and yields the largest accuracy gains when global sampling policies are suboptimal (i.e., at low 𝛽_c_values).

In sum, RNNs successfully acquired the directed-exploration policies when trained with appropriate objectives, but failed to spontaneously produce early streaking. This dissociation underscores that streaking is not a default outcome of optimizing task performance in standard recurrent architectures. Instead, it may reflect specific inductive biases or cognitive constraints in human learners not captured by these models. We next examined its functional consequences for human performance and its relationship to individual cognitive traits.

### Early streaking co-varies with uncertainty-guided sampling across participants

Using standard RNNs as a computational benchmark, we observed a key dissociation: the networks successfully learned global, near-optimal uncertainty-guided sampling but failed to acquire early streaking. This dissociation raises a critical question about human behavior: given that humans exhibit the dual policy at the group level, how do individuals balance these two components? One theoretical perspective would predict a tradeoff: individuals who rely more heavily on early streaking might show weaker uncertainty-guided sampling, as if allocating cognitive resources toward one strategy comes at the expense of the other. Alternatively, both strategies could reflect a coordinated dual policy for epistemic exploration. Under this view, individuals who engage in more pronounced early streaking may also demonstrate stronger uncertainty-guided sampling, with both components contributing synergistically to decision performance.

To test these predictions, we compared participants whose behavior was best explained by models with versus without a streaking mechanism. We performed model selection using the evidence lower bound (ELBO), which balances model fit against complexity^27,29^. In the MATCH condition, the model including an initial streaking phase was far less prevalent than the simpler model without it (log_10_BF_10_ = -106.9, 95% CI [-150.7, -62.5]; model with fixed *θ* at 0: P_exc_ > 0.99). By contrast, in the GUESS condition, the inclusion of an initial streaking phase markedly improved model fits for most participants (286 out of 420 participants; log_10_BF_10_ = 753.4, 95% CI [643.4, 866.5]; model with free *θ*: P_exc_ > 0.99; Fig. 7A; see also Supplementary Fig. S12A).

**Figure 7.**
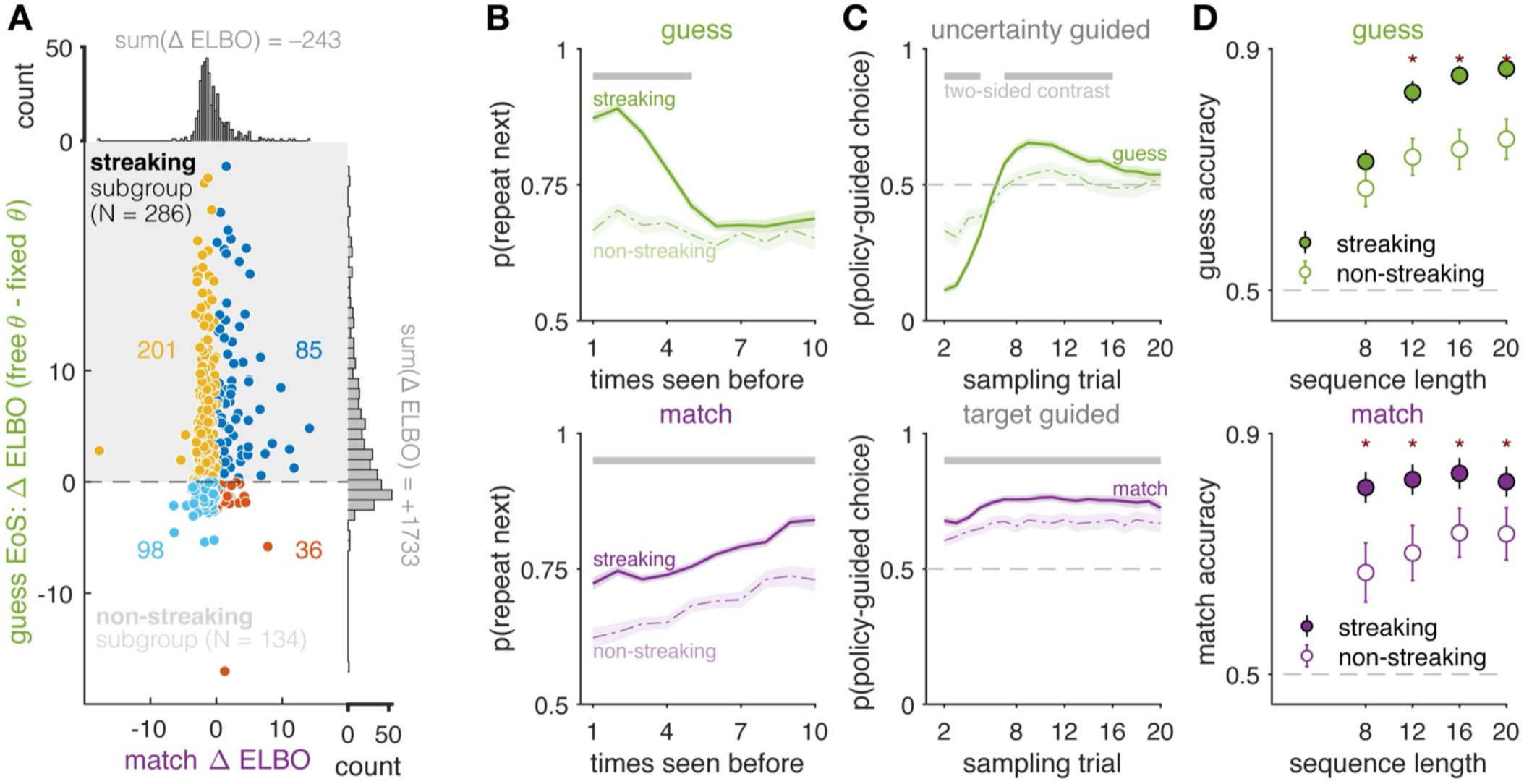
Distinct behaviors by early-streaking tendency. Top row: GUESS condition. Bottom row: MATCH condition. (**A**) Participants grouped by the ELBO difference between a model with a free threshold *θ* and a model without an initial threshold (*θ* fixed to 0). Streaking subgroup classification was based on the ELBO difference in the GUESS condition, where streaking is not confounded with reward-driven choice repetition. See Supplementary Fig. S13 for grouping based on ELBO in the MATCH condition, for completeness. (**B**, **C**) Fraction of choices consistent with policy-guided sampling, shown separately for participants grouped by early-streaking tendency. In the MATCH condition, “policy-guided” means target guided; in the GUESS condition, it means uncertainty guided (definitions as in Fig. 2B). (**D**) Choice accuracy for groups defined by model evidence favoring presence or absence of an initial sampling threshold. Accuracy as in Fig. 2A (MATCH: target-choice rate; GUESS: final-probe correctness).

We divided participants into two groups based on model fits in the GUESS condition: those better explained by a model with an initial evidence threshold (“streaking” subgroup: *N* = 286) and those better explained without it (“non-streaking”: *N* = 134). By definition, the streaking participants showed positive *Evidence of Streaking* (EoS), while non-streaking subgroup showed negative EoS (Fig. 7A; Supplementary Fig. S12A).

As expected, streaking participants were more likely to resample options in continuous streaks during the early trials of the GUESS game (Fig. 7B). Later in the sequence, they also made better use of uncertainty-guided sampling strategy, scheduling actions that best reduced uncertainty about the hidden state (Fig. 7C). In the MATCH condition, streaking participants showed a stronger preference for the target-colored option, which is the optimal strategy there (Fig. 7C). Across both conditions, they also repeated their previous choice more often, although this bias did not extend to stronger initial repetition in the MATCH game (Fig. 7B, lower panel). The streaking subgroup had lower leak and noise than non-streaking subgroup, but noise remained a nontrivial source of suboptimality in the streaking participants: the noisy model was decisively favored in both conditions (expected posterior family frequency > 0.98; Supplementary Fig. S12B–C).

This pattern reveals a mixture of strategies. Streaking participants combined an early suboptimal tendency to repeat choices with later, more adaptive directed-exploration policy. Moreover, this combination was linked to higher performance in both conditions (MATCH: t(217.0) = 6.13, p < 0.0001, d = 0.73; GUESS: t(204.4) = 7.61, p < 0.0001, d = 0.91; Welch’s t tests, Fig. 7D). Thus, local streaking was not a sign of disengagement or inattention, but part of a broader strategy that, overall, improved performance.

Taken together, our results suggest that human participants combine optimal and suboptimal policies to serve epistemic exploration. Moreover, despite being suboptimal from a normative perspective, early streaking improved accuracy in the GUESS condition. By contrast, RNNs optimized for hidden-state inference and meta-cognitive uncertainty reduction did not develop this behavior despite extensive training. In this setting, streaking appears to be a distinctive feature of human information gathering: an apparently suboptimal bias that was not reproduced by our optimization benchmarks and that nonetheless enhances inference performance.

### Early streaking and uncertainty-guided sampling are related to distinct psychological traits

In light of the wide variability in sampling patterns across participants (for example, streaking vs. non-streaking), we asked which psychological traits were related to these individual differences. We first examined how individual sampling policies related to performance in the GUESS condition. Streaking is a suboptimal strategy from a normative standpoint, yet it covaried with greater reliance on uncertainty-guided choices. To quantify streaking tendency as a continuous model-based index, we used model evidence EoS favoring the early streaking policy rather than a binary grouping.

Regression analyses showed that EoS contributed positively to GUESS accuracy over and above *β*_c_ and independently of the detrimental influences of leak *δ* and learning noise *σ* (Fig. 8A, left). Nested likelihood-ratio tests indicated that removing EoS significantly worsened regression model fit, χ²(1) = 8.31, p = 0.004, and removing *β*_c_ also worsened fit, χ²(1) = 20.84, p < 0.001, indicating that both EoS and *β*_c_ make unique, positive contributions to GUESS accuracy.

**Figure 8.**
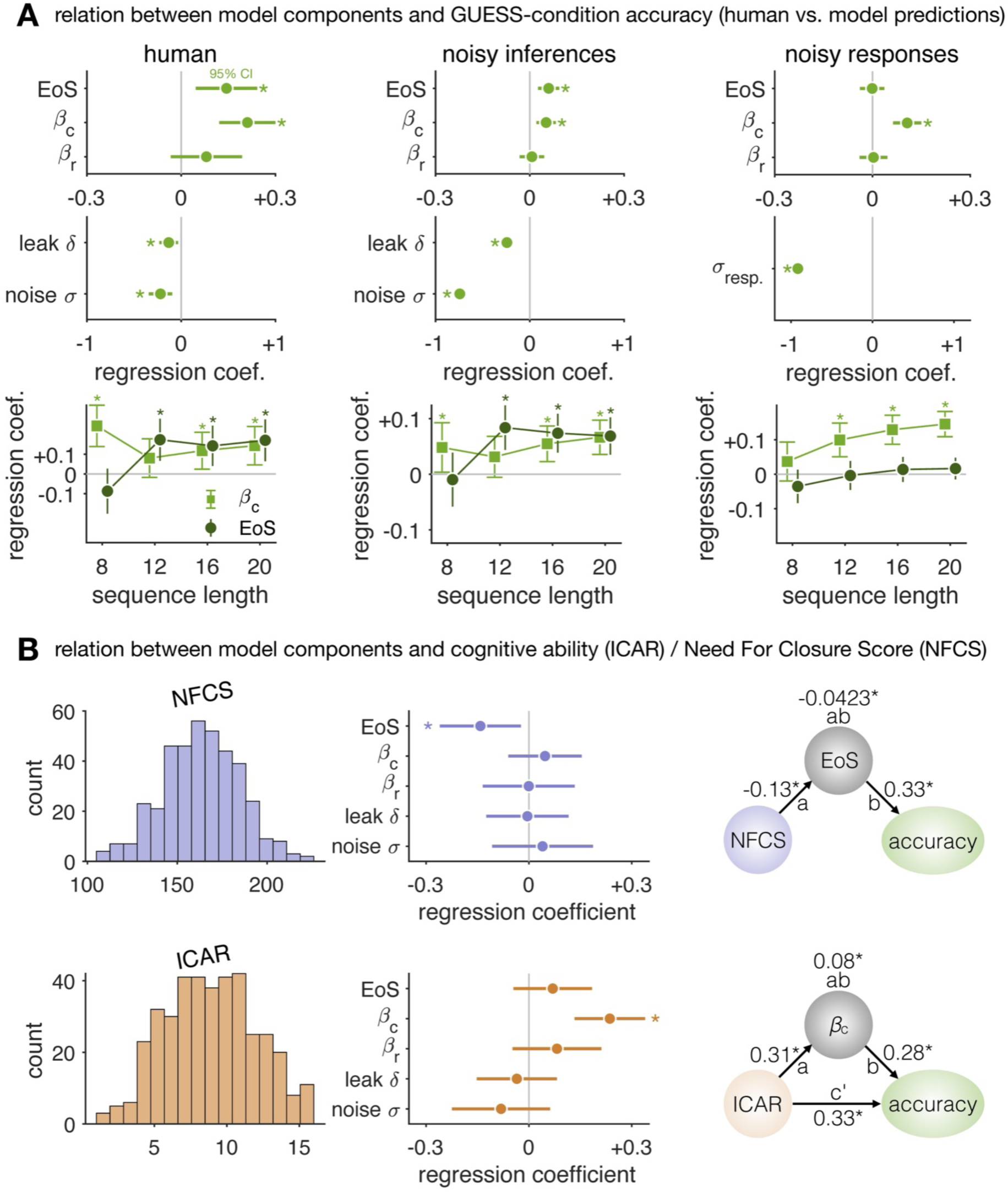
Relationships between performance, policy, cognitive ability, and psychological traits. (**A**) Inference accuracy in the GUESS condition was jointly predicted by *β*_c_ (uncertainty-directed exploration) and EoS (early streaking-based information seeking), controlling for other model parameters. Ex post simulations of a leak + noise observer (“noisy inferences”) reproduced these regression effects on model-predicted accuracy. In a control model with response noise (*σ*_resp._) at the time of the final probe but no learning noise during sampling, only *β*_c_ positively predicted simulated accuracy, while EoS was not significant. Top panels: mean coefficient is shown by the circle; 95% CI by the green horizontal bar; asterisks indicate significant effects against zero (two-sided one-sample t-tests). Bottom panels: regression analysis individually for each sequence length. (**B**) Left: histograms of questionnaire scores; middle: hypothesis-driven tests of selective associations (circle: mean; horizontal bar: 95% CI); right: mediation analyses (also see Supplementary Tables 1–2).

The benefit of early streaking was most pronounced in late trials of longer sequences, that is, more than 12 choices. Using accuracy predicted by the full leaky-noisy model as the dependent variable reproduced the same pattern (Fig. 8A, middle; Supplementary Table 1). By contrast, in an exact model without learning noise *during* sampling, where noise was applied only as response variability at the final guess *after* sampling, EoS no longer improved accuracy and only *β*_c_ predicted performance (Fig. 8A, right; Supplementary Fig. S14). Eliminating learning noise, but not leak, eliminated the streaking benefit on simulated accuracy (Supplementary Fig. S15; see also Supplementary Fig. S16, where higher learning noise spreads the benefit across more trials). Together, these results indicate that streaking reinstates structured sampling and enhances expected accuracy of inference, particularly in the presence of noise during belief updating.

Given the central role of EoS and *β*_c_ in supporting better inference accuracy, we asked how these components relate to broader dispositions. We used a top-down framework with a priori hypotheses for two established scales collected in our online sample (Fig. 8B): The Need for Cognitive Closure Scale (NFCS) and the International Cognitive Ability Resource (ICAR). NFCS indexes preference for a rapid conclusion, whereas ICAR indexes general cognitive ability. We hypothesized that NFCS would relate (negatively) to EoS (willingness to continue sampling before reaching a conclusion), and that ICAR would relate (positively) to *β*_c_. Specifically, individuals with high need for closure seek rapid conclusions and may sample less extensively, whereas those with lower NFCS may be more comfortable with prolonged, structured sampling. And individuals with higher ICAR scores may engage in more uncertainty-guided exploration, which requires accurately tracking which option is more uncertain.

NFCS and ICAR were uncorrelated in our sample (bootstrap mean Spearman *ρ* = -0.06, p = 0.2; Supplementary Fig. S17), indicating minimal shared variance. Multiple regression confirmed a double dissociation: only EoS significantly predicted NFCS (*β* = -0.142, 95% CI [-0.260, -0.023], t(381) = -2.36, p = 0.019), and only *β*_c_ significantly predicted ICAR, *β* = 0.236, 95% CI [0.133, 0.339], t(385) = 4.51, p < 0.0001 (Fig. 8B; Supplementary Table 1). This dissociation indicates that motivational trait (NFCS) and cognitive ability (ICAR) shape distinct computational substrates of inference.

Mediation analyses further clarified the relations linking traits to policies to performance. NFCS did not directly predict accuracy (c’ path: p = 0.921), but only indirectly through its negative relation to EoS (indirect path: a*b = -0.0423, 95% CI [-0.0805, -0.0052], bootstrap p = 0.0236; Fig. 8B): higher NFCS scores predicted lower EoS, which in turn reduced accuracy (Supplementary Table 2). This suggests that the drive for rapid closure works against the patience required for streak-based sampling of each novel option in turn, even though we have seen that such structured sampling improves performance.

By contrast, ICAR predicted accuracy both directly and indirectly through *β*_c_. Participants with higher ICAR scores showed stronger sensitivity to uncertainty, and both ICAR and *β*_c_ independently predicted higher accuracy. *β*_c_ also predicted ICAR, consistent with a reciprocal association in which cognitive ability supports adaptive inference strategies and strategy use reflects general ability. Together, these findings indicate a layered architecture in which stable traits shape task performance through specific latent computational processes.

Finally, in a complementary bottom-up analysis, we tested whether personality measures predicted individual differences in sampling policies. Guided by evidence that many standalone traits can be situated within the Big Five taxonomy^30^, we used leave-one-participant-out cross-validation to predict NFCS, ICAR, *β*_c_, and EoS from Big Five scores^31^. NFCS was indeed predictable (R² = 0.32; Supplementary Fig. S18; Supplementary Table 3), with positive weights for Conscientiousness and Negative Emotionality and negative weights for Open-mindedness and Extraversion. By contrast, ICAR, *β*_c_, and EoS were only weakly predicted (R² ≤ 0.03), indicating that these constructs were not well captured by broad personality dimensions. ICAR was better explained by the model parameters (R² = 0.11; see Supplementary Fig. S18–S19), consistent with the view that general cognitive ability reflects the efficiency of latent task-specific processes, while NFCS maps onto stable personality axes. This dissociation highlights the value of computational modeling: whereas psychological traits such as need for closure can be subsumed within established dispositional taxonomies, task-specific cognitive policies such as streaking and uncertainty-guided exploration reflect distinct latent computational phenotypes that are not reducible to personality structure.

## Discussion

We examined human information seeking when prospective valuation was removed (our novel GUESS condition) and compared it with reward-driven sampling (MATCH condition, resembling a classic two-armed bandit). Human behavior revealed a dual structure of information seeking: a global policy of uncertainty-guided sampling and a local policy of early streaking. Globally, choices were guided by reward in the MATCH condition and by uncertainty in the GUESS condition. Locally, upon first encountering a novel option, observers resampled it several times in a streak before switching, consistent with testing a provisional hypothesis to establish a satisficing level of certainty rapidly. When prospective valuation in payoff units was reinstated, as in the MATCH condition, this dual-policy structure disappeared. Across individuals, stronger streaking was associated with lower need for cognitive closure and with higher decision accuracy under noisy belief updating. These findings reconcile human suboptimality with adaptiveness, showing how a structured, local policy such as streaking can improve inference accuracy under imprecise belief updating.

### Information seeking versus reward seeking

Neuroscience research presents contrasting views on how the brain values information. One framework treats information as intrinsically rewarding: information gain and uncertainty reduction may serve as internal reward signals^32^. Supporting this view, midbrain dopamine neurons encode non-instrumental information—cues that predict upcoming rewards without altering their probability—in ways that mirror reward prediction^13,33^. Orbitofrontal cortex similarly represents information value alongside reward^15,34^. This implies a common neural currency for evaluating both information and external rewards. However, other work emphasizes that information seeking can be guided by motives beyond reward seeking, including anticipated affective and cognitive consequences of knowledge^12,18,35–37^. A central challenge has been dissociating information value from reward statistics, as information inherently facilitates reward learning, and reward sustains task motivation in most experimental paradigms. Our design addresses this by holding visual statistics constant across conditions while manipulating only the role of outcomes: in the GUESS condition, samples revealed a latent category rule instrumental for final accuracy but not expressible in reward units during sampling, whereas in the MATCH condition, the same samples carried immediate payoff. This controlled contrast isolated goal-driven shifts in exploration policy, revealing that epistemic information seeking deploys distinct computational strategies beyond what reward-based frameworks predict, allowing for future delineation of neural underpinnings.

### The computational underpinnings of early streaking

Why do participants concentrate early sampling on a single option? We propose that early streaking reflects a divide-and-conquer logic shaped by several cognitive constraints. First, by focusing on one option initially, the cognitive system reduces switching costs, e.g., the metabolic and computational expense of shifting attention and updating multiple belief states^38^. This is consistent with the general repetition bias (*β*_r_ > 0) we observed across both task conditions, indicating a default tendency to avoid switches.

Second, this focused sampling stabilizes state representations and sharpens coding precision for the currently active hypothesis before a switch occurs^39,40^. Early repetitive sampling may thus reflect a cognitive schema in which participants generate a provisional hypothesis about each option before testing it through subsequent choices. Such hypothesis-testing strategies have been argued to improve efficiency in self-directed learning, since they deliver evidence perceived as more relevant at each choice^41^. Importantly, streaking should not be mistaken for simple stickiness or entrapment in a policy “fixed point.” Here we observe a distinctive pattern in active sampling behavior: participants continued to resample until evidence confirmed their current belief. They did not switch away in the face of ambiguity, but only once confirmatory evidence had accumulated (Supplementary Fig. S1B), underscoring the purposeful, confirmatory nature of the streak.

Alternatively, could streaking be attributable to asymmetric memory decay? Decay refers to the loss of accumulated evidence when an option is left unchosen. If streaking simply reflected selective maintenance, one would expect higher leak for unchosen options. Instead, the best-fitting model indicated a common leak across options, which provided a decisively better description of behavior than models allowing asymmetric decay. Moreover, estimated leak was lower in the GUESS condition than in the MATCH condition, yet streaking was stronger in GUESS. This dissociation rules out memory decay as the driver of streaking.

Could delayed rewards explain streaking? In a control “FIND” condition in our previous work^21^, participants pursued a delayed reward (identifying a target color) but were informed of the goal *before* sampling. This allowed for prospective outcome valuation, similar to typical Decisions From Experience paradigms^7^. The results showed a clear dissociation: “FIND” significantly diminished streaking compared to the purely epistemic (GUESS) condition, yet it still yielded a significant increase in uncertainty-guided exploration relative to the immediate-reward (MATCH) condition. This strongly suggests the mechanism of streaking is not delayed reward, but the release from prospective outcome valuation during sampling.

### Adaptive function of early streaking in the presence of noisy belief updating

In human participants and in RNNs, uncertainty-guided sampling unsurprisingly improved inference performance. In our human participants, however, streaking also benefited inference performance, and most strongly so under noisy belief updating. By rapidly accumulating evidence on one option, the policy limits the dilution of updates by random fluctuations in imprecise learning^23,24,42^, thereby stabilizing beliefs^43^. In this sense, streaking illustrates how a policy that may appear locally suboptimal can be adaptive in the presence of irreducible neural noise, consistent with accounts in which structured, statistically suboptimal strategies can improve decision quality^44,45^. From a neural-coding perspective, streaking improves precision early when beliefs are weak. Neural coding precision of belief-evidence consistency is poor at low belief magnitude^46^, so an early sequence of choice such as AAAAA should yield clearer consistency decoding than an interleaved ABABA sequence at matched trial counts.

### Dissociation between global and local policies in recurrent neural networks

Our RNNs did not acquire early streaking, even though they successfully learned the optimal policy of uncertainty-guided sampling when trained to maximize belief certainty. This dissociation is theoretically informative: it demonstrates that RNNs possess the computational capacity to approximate near-optimal exploration policies yet do not spontaneously acquire the local streaking rule that humans deploy.

We do not claim that streaking is impossible for any artificial network architecture. Rather, we use RNNs as a controlled computational benchmark precisely because we can systematically manipulate their optimization objective (i.e., the goal they pursue) while holding other factors constant. Rather than treating networks as black boxes, we use them to test which objectives are causally necessary for particular cognitive strategies to emerge. This mirrors our experimental manipulation in humans, where identical visual statistics yielded different exploration policies depending solely on task goals. The critical finding is the selective dissociation: RNNs trained on our task acquired one aspect of human participants’ dual information-seeking policy (i.e., uncertainty-guided sampling) but not the other aspect (early streaking).

Streaking bears resemblance to curriculum learning effects, particularly the “blocking” advantage. Humans often exhibit a blocking advantage, learning better when related information is presented in contiguous blocks that make the underlying structure more salient. This phenomenon has been extensively studied in the curriculum learning literature^47–49^. While biologically inspired gating networks can reproduce a blocking advantage when the curriculum is externally imposed^48,50–53^, these models do not actively control their own sampling. The blocking benefit in these cases arises from how the imposed input structure interacts with the network’s dynamics, rather than from an emergent sampling policy chosen by the agent itself. Future work should investigate whether architectures endowed with mechanisms for self-directed sampling can discover streaking policies autonomously, thereby bridging the gap between externally imposed curriculum effects and intrinsically generated sampling policies.

### Individual differences in epistemic sampling

Individual differences in exploratory tendencies have been linked to broader behavioral and cognitive features^54–56^, suggesting a phenotypic signature of adaptation to environmental and evolutionary pressures^57^. Foundational work has shown that such differences are stable over time and relate to mental health outcomes^58^, motivating systematic characterization of how individuals vary in information-seeking strategies. We framed individual differences at three linked levels: person-level dispositions, latent processes, and task policies. Traits such as need for closure (NFCS) and abilities measured by the ICAR scores index broad dispositions and capacities; Latent processes center on noisy belief updating; Policies are the strategies expressed in the task. Our results uncover a dual policy in epistemic information seeking: more pronounced streaking (higher EoS) accompanies stronger uncertainty-guided sampling (higher *β*_c_) in the same individuals, and both independently improve decision performance. Crucially, the two policies map onto distinct dispositions. *β*_c_ indexes a global policy that allocates samples by uncertainty and correlates positively with ICAR scores, consistent with a domain-general capacity that supports meta-cognitive use of uncertainty. By contrast, EoS relates to lower NFCS and indexes a local, bounded policy that builds rapid certainty under noisy belief updating through early streaking. This pattern mirrors the dissociation between humans and RNNs, where a meta-policy RNN learned uncertainty-guided sampling yet did not develop streaking.

### Conclusions

Our results reveal a dual policy architecture combining global directed-exploration with local streaking. This dual policy emerges when reward pricing is absent, but shifts to reward-guided sampling without streaking when expected-payoff evaluation is present. Streaking highlights how stable traits may not directly map to performance in cognitive tasks, but instead shape the policies people deploy to gather information. This layered view pushes the field beyond the search for “trait correlates of exploration” and toward an understanding of how person-level dispositions tune the latent processes that govern decision policies.

## Methods

### Participants

The target sample size was set to 525 based on both empirical considerations and a priori power analyses. Based on our existing laboratory dataset^21^, an a priori power analysis indicated that 186 participants were required to detect the smallest condition-wise contrast observed previously (repetition hardness *ε*; d_z_ = 0.183) with 80% power at a one-sided 5% significance level. No prior study had directly examined the relationship between personality measures and this task, so we set the minimal effect of interest to a correlation of *r* = 0.15. Detecting such an effect with α = 0.05 (two-tailed) and 80% power requires 392 participants. Accounting for the typical attrition rate in online studies (∼30%), we set the recruitment target at 509 participants.

Participants were recruited via the Prolific platform (www.prolific.co) and were informed about the purpose and content of the study both on the platform and immediately prior to the experiment. The study was approved by the INSERM Ethical Review Committee (IRB00003888). All participants provided informed consent electronically before participating. They were compensated £8 for approximately 60 minutes of participation, with an additional £2 bonus contingent on good task performance. Participants were informed that the bonus would be awarded if they earned enough medals (see below); 79 participants met this criterion and received the bonus.

#### Inclusion/exclusion criteria

Eligible participants were required to be UK residents, fluent in English, between 18 and 35 years of age, with normal or corrected-to-normal vision (no color blindness), and to have completed at least five previous Prolific studies with an approval rate of ≥ 80%. In addition, they must not have taken part in previous laboratory studies using similar colored stimuli. To ensure data quality, we applied several exclusion criteria. For data analyses, 105 participants were excluded, resulting in a final dataset of 420 participants.

Exclusion criteria were: (1) incomplete task performance; (2) missing trials on more than five games due to technical issues; (3) production of stereotyped choice patterns (> 50% across all conditions), defined as choice sequences where participants either chose only one option throughout, only one response side, or systematically alternated between options or sides across the entire sequence, suggesting blind responding without outcome consideration, or (4) failure on task comprehension checks (the first four items of the debriefing questionnaire).

The retained participants’ performance was comparable to that observed in the laboratory^21^, indicating the reliability of the online data. For the analyses linking task behavior with questionnaire responses, we further excluded 28 participants who either failed to complete the questionnaires or responded incorrectly to at least one of two catch items inserted in the NFCS and BFI-II questionnaires. In addition, one participant was excluded because all four parameter estimates (*β*_c_, *β*_r_, leak *δ*, noise *σ*) from the computational model fit to the GUESS condition were identified as outliers using Tukey’s fences (interquartile range method). The final dataset used for the combined behavioral and questionnaire analyses comprised 391 participants (mean age = 27.0 years, SD = 4.8; 190 female). For NFCS regression against model parameters, four influential outliers were additionally identified via regression diagnostics. Results were robust to their exclusion: while model fit improved (log-likelihood increased by 19.99; AIC decreased from 1115 to 1075), all parameter estimates remained qualitatively unchanged (Supplementary Table 1).

### Task and experimental procedure

#### Procedure overview

The study comprised two phases: (1) the main experiment and (2) the questionnaires. Participants first received general information about the study and its procedure. After providing informed consent, they performed the main behavioral task, which consisted of (a) a step-by-step tutorial explaining the rules, (b) eight practice game sequences, and (c) four blocks of 16 games each, forming the main task (64 games in total, 896 sampling trials). Participants were encouraged to take self-paced breaks between blocks, but not within blocks. After the task, they could take another break before beginning the questionnaire phase, which included (a) a short debriefing on the task, (b) the Need-for-Cognitive-Closure-Scale (NFCS) questionnaire^59,60^, (c) the 60-item Big Five Inventory II (BFI-II)^31^, and (d) a subset of the International Cognitive Ability Resource (ICAR) items^61,62^. Finally, participants were informed whether they had earned the £2 bonus and were invited to provide optional feedback.

#### Task design

The experimental task framing followed our previous foundational study, which first established an epistemic task condition untied from prospective reward pricing^21^. Participants were instructed that they would be sampling gems from two candidate “bags.” Each bag contained only one type of gem of a specific shape (e.g., one bag contained triangles, the other circles), but the individual gems could vary in color shade. On average, the blue bag contained predominantly more bluish shades and the orange bag more orangish shades, though each still probabilistically included about one-third of samples in the non-dominant color (Fig. 1B).

Each sampling sequence (a “game”) involved two new bags. At the start, participants did not know the underlying color distributions linked to the bags and had to sample sequentially, one gem per trial, from either bag. They observed only the sampled gem’s color shade, not the outcome of the unchosen bag. Sampling sequences ended unpredictably, after 8, 12, 16, or 20 trials with equal chance. Two types of games (conditions) were randomly interleaved:

- *MATCH game*. The goal was to sample gems with the most intense shades matching a target color (blue or orange, counterbalanced across game sequences). Each choice contributed directly to the participant’s performance score.
- *GUESS game*. Participants were instructed to “discover” the bags’ colors during sampling and, at the end of the sequence, to “guess” the dominant color of one randomly selected bag in a 2AFC question (“orange or blue?”). Here only the accuracy of this final inference counted toward the performance score.

Thus, the MATCH game emphasized immediate reward maximization, whereas the GUESS game required information seeking about the hidden structure. The structure was the mapping from the bag (and its specific gem shape) to its dominant gem color. Participants were clearly informed at the start of each sequence which game type they were playing. They were also told that (i) the sequence could end at any time, (ii) the dominant color of one bag could not be inferred from that of the other bag (both could be blue, both be orange, or one of each), and (iii) even in a blue bag, some orange gems could appear, though these tended to be less intense.

#### Trial structure

Each game began with an instruction screen presenting the two new bags (and their gem shapes) and specifying whether the game was a MATCH or a GUESS condition. Pressing the “space” bar initiated the sampling trials. On each trial, the two gem shapes appeared randomly on the left or right side of the screen, to be selected using the “f” and “j” keys, respectively. Each trial was comprised of:

1. a probe screen displaying both gem shapes until a choice was made;
2. an outcome screen showing the sampled gem shape in its color shade for 500 ms;
3. an inter-trial interval of 250–500 ms.

In GUESS games, sequences ended with a 300 ms blank screen, followed by a final question screen displaying one of the gems and two response buttons (orange and blue, randomly assigned to the left/right sides). Participants judged the probed bag’s dominant color, saw their selection highlighted (300 ms), and received feedback (the true color for 500 ms). The game then ended with a 300 ms blank screen before the next instruction screen.

#### Feedback and incentives

The 64 games were divided into four blocks of 16, with equal numbers of MATCH and GUESS conditions. At the end of each block, participants received feedback in the form of medals (none, bronze, silver, gold) based on their average performance score: (1) < 0.50: no medal (0 points); (2) 0.50–0.65: bronze (1 point); (3) 0.65–0.70: silver (2 points); (4) ≥ 0.70: gold (3 points). Participants earned the £2 bonus if they accumulated at least 6 points across the four blocks. The score was the average of game-level scores in a block: for GUESS condition, it was the mean accuracy of the final questions; for MATCH condition, it was the fraction of choices that matched the predictions of a near-optimal reward-maximizing model with a small learning noise of 0.5. Thus, in the MATCH condition, the incentives encouraged prospective valuation of individual sample outcomes.

#### Stimuli

Eight geometric shapes were combined into 28 possible gem-shape pairs (Fig. 1B). Each hidden bag was associated with a color distribution (2/3 dominant, 1/3 non-dominant). Color shades were chosen such that the log-evidence for the underlying distribution scaled linearly with color intensity, ensuring that each outcome carried information gain regardless of whether the dominant color was blue or orange. Outcome sequences were pre-generated, equating theoretical difficulty across conditions by simulating artificial agents.

The same 32 sequences were used across the two conditions (i.e., direct replica), with only the instructions differing between MATCH and GUESS. The 64 total sequences were arranged into four pseudo-randomized blocks of 16, balancing condition type, bag–color mappings (blue/orange, both blue, both orange), and gem-shape repetition across consecutive games.

#### Questionnaires

After completing the main task, participants answered a set of debriefing questions and then completed two self-report questionnaires and a reasoning test, used as a proxy for IQ. The order was fixed across all participants: (1) Debriefing questions. Participants first responded to seven questions assessing their understanding of the task instructions. (2) Need for Closure Scale (NFCS)^59,60^. The NFCS is a 41-item questionnaire measuring individuals’ “need to settle for any answer, as opposed to further sustaining ambiguity.”^60^ Scores up to 82 indicate a low need for closure, while scores above 205 indicate a high need for closure^60^. (3) Big Five Inventory II (BFI-II)^31^. The BFI-II is a 60-item questionnaire assessing the five broad domains of personality: Extraversion, Agreeableness, Conscientiousness, Negative Emotionality (formerly Neuroticism), and Open-Mindedness (formerly Openness to Experience). Each domain can be further divided into three facets (15 in total). (4) Reasoning test. We administered a 16-item subset of the International Cognitive Ability Resource (ICAR)^61,62^, including 4 matrix reasoning items, 4 three-dimensional rotation, 4 letter series, and 4 verbal reasoning items. Each item was multiple-choice. This subset has been validated in previous work^63^ and was used here as a measure of general cognitive ability. Scoring was one point per correct response.

### Computational model of sequential sampling

The probability of generating a gem of a given color shade 𝑐 from the orange bag was defined by a sigmoid function:

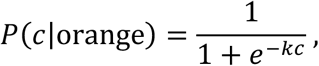

with 𝑐 ∈ [−1, +1] (-1 = pure blue, +1 = pure orange) and slope 𝑘 > 0 (Fig. 1B). The complementary probability was 𝑃(𝑐|blue) = 1 − 𝑃(𝑐|orange). The corresponding log-likelihood ratio is logLR = 𝑘𝑐, representing the Bayesian log-evidence (𝑒) in favor of the orange- over blue-bag hypothesis (Fig. 1B). This means that the color shades directly convey evidence about the hidden state, making the task both interpretable and tractable for modeling. This mapping from color intensity to log-likelihood ratios and the mixture distributions replicate the specifications introduced by Alméras et al.^21^ The slope parameter was set to 𝑘 = 1.4386, yielding an error rate of roughly one third for observing a bluish outcome from the orange bag (and vice versa).

The computational model of sampling behavior included a learning module identical across both conditions. At each trial 𝑡, the model accumulated evidence *𝑒_i,t_* for the chosen option 𝑖 to form an evolving posterior belief 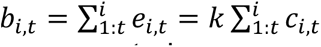. The unchosen option received no update. A utility variable was then computed as:

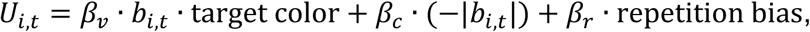

where 𝛽*_v_* (MATCH condition only), 𝛽_c_, and 𝛽*_r_* are sensitivity parameters to the target evidence, belief uncertainty (−|𝑏_i,t_|), and repetition bias, respectively. The “target color” was coded as −1 when the goal in the MATCH condition was to choose the gem most likely drawn from the blue bag, +1 when the target was orange, and 0 in the GUESS condition. The repetition bias was coded as +1 if the previous choice at trial 𝑡 − 1 was option 𝑖, and -1 otherwise, thus biasing choice to repeat on trial 𝑡 when 𝛽_r_ > 0.

Choices were generated by one of two policies. By default, a softmax rule was applied to the utilities, with the 𝛽 parameters jointly determining the effective inverse temperature,

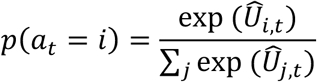

In addition, the model allowed for an initial sampling phase: as long as the posterior belief for the chosen option remained below a threshold level 𝜃, choices ignored utility and followed a sticky rule, repeating the previous choice with probability 𝜀 while switching with probability 1 − 𝜀. This phase was thus governed by 𝜃, which sets the amount of posterior belief required before switching to utility-based choice policy, and by 𝜀, which determines the strength (hardness) of repetition under the threshold 𝜃. To avoid discontinuities in the likelihood, 𝜃 was modeled as normally distributed with a small standard deviation of 0.1 in logLR units (notably small relative to the information provided by a single sample, up to ∼1.44 logLR units).

Note that the computational model nests the optimal sampling agent for both conditions. The optimal agent does not include the initial sampling phase or the repetition bias. In the MATCH condition, it selects based solely on the signed difference between the mean color-mixture estimates relative to the target color, with 𝛽_)_ → +∞ (argmax choice). In the GUESS condition, it selects based solely on the difference in accumulated information between options, with 𝛽_!_→ +∞.

Following each outcome, the posterior belief about the chosen option was updated and further corrupted by additive random learning noise drawn from a normal distribution of zero mean and standard deviation 𝜎:

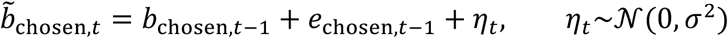

Moreover, a memory leak caused beliefs to drift back toward zero between trials:

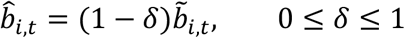

with the leak parameter 𝛿 shared across chosen and unchosen options. Model comparison indicated that this common-leak model provided a better fit to participants’ choices than a model alternative with independent leaks for chosen and unchosen options (Supplementary Fig. S7). The final noisy-leaky utility used for choice selection was then:

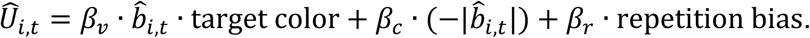

### Computational model fitting

We estimate d the conditional likelihood of each participant’s decisions given the model parameters using Sequential Monte Carlo methods (particle filtering). Point estimates of best-fitting parameter values were first obtained via Bayesian Adaptive Direct Search (BADS)^64^, using 10 random starting points per parameter. These point estimates then served as initializations for estimating the joint posterior distribution via Variational Bayes Monte Carlo (VBMC)^27,29^. Parameter priors used in the VBMC fitting are available inside the model-fitting script in the online repository. Reported best-fitting parameters correspond to the posterior means estimated by VBMC, and model goodness-of-fit was quantified using the evidence lower bound (ELBO).

### Ex post simulations of best-fitting computational model

We simulated the models in generative mode using their best-fitting parameter values (Fig. 4A; Fig. 5C, F). This procedure assessed whether, given the retrieved parameters, the models could reproduce the behavioral effects of interest without replaying participants’ exact choice sequences and outcome history. For the GUESS condition, we evaluated out-of-sample inference by simulating each game and extracting the model’s final posterior belief, after the last update of the chosen option. We then applied an argmax rule to obtain the categorical response (orange if the posterior belief > 0, blue if < 0). GUESS accuracy was scored by whether this response matched the ground-truth bag color (hidden state), with chance-level performance coded as 0.5 when the posterior belief was exactly zero. Because model parameters were estimated from sampling behavior only, these predictions of GUESS accuracy were therefore “out-of-sample” with respect to the fitted domain.

### Model comparison

We performed random-effects Bayesian model selection (BMS) using the standard Dirichlet parameterization^65^. Fixed-effects comparisons were conducted using Bayes factors, by default contrasting more complex models with simpler ones (e.g., model with a free threshold vs. model without a threshold; Supplementary Fig. S12A). For each condition, the ELBO across participants was entered into BMS to obtain exceedance probabilities along with Dirichlet-based posterior probabilities and 95% confidence intervals. In addition, we ran factorial family-wise model comparisons (e.g., “leak” vs. “no leak” and “noise” vs. “no noise”; Fig. 5B, E), based on grouping models into families and estimating family exceedance probabilities under the random-effects assumption. Confidence intervals for family probabilities were computed by Dirichlet sampling (10,000 draws).

### Model recovery

We conducted model recovery to verify that the models under comparison were both distinguishable and recoverable within the fitting procedure. Using the mean best-fitting parameters from the full model across all participants and both conditions (thus avoiding condition-specific bias; except with 𝛽*_v_* = 0 in the GUESS condition), we simulated data and then fit all four model variants (full model, or knocking out leak, noise, or both) to these simulations. Subsequent model comparison confirmed that each model was both identifiable and recoverable (Supplementary Fig. S5).

### Model parameter recovery

We performed model parameter recovery to test whether our fitting procedure introduced spurious parameter dependencies. Simulated choices were generated either from participants’ best-fitting parameters (Supplementary Fig. S6A) or from random draws within predefined uniform ranges (*δ* ∈ [0, 0.5], *σ* ∈ [0, 4], *β*_v_, *β*_c_, *β*_r_ ∈ [−3, 3], *θ* ∈ [0, 4], *ε* ∈ [0.5, 1]; Supplementary Fig. S6B). Correlations between generative parameters were destroyed by random permutation. Refitting the model showed no spurious correlations and the parameters themselves were recoverable (Supplementary Fig. S6C).

### Analysis of information sampling patterns

#### Policy-aligned choice

On each trial t, we computed each option’s running cumulative log-evidence from trials 1 to t − 1. We then identified (i) the option most consistent with the target in the MATCH condition, defined as the option with the largest accumulated log-evidence toward the target color, and (ii) the most uncertain option in both conditions, defined as the option with the smallest absolute accumulated log-evidence. A choice on trial t was coded as aligned with the target-guided policy if it matched the option most consistent with the target (applicable to the MATCH condition only), and as aligned with the uncertainty-guided policy if it matched the most uncertain option. Each indicator equals 1 if aligned and 0 otherwise. In the event of ties, the trial was excluded for that policy.

#### Signed evidence

For the GUESS condition, we compared the sign of the outcome on trial t with the sign of the prior belief, given by the accumulated log-evidence from trials 1 to t − 1. By task convention, signs were > 0 for orange and < 0 for blue. Outcomes with the same sign as the prior belief were coded as consistent and outcomes with the opposite sign as conflicting. For the MATCH condition, the outcome on trial t was coded as aligned if it matched the sign of the target color and misaligned otherwise.

### Neural network architecture

The artificial networks for the MATCH and GUESS conditions were standard Elman RNNs. At each trial, the input 𝑋*_t_* (color shades ∈ [−1, 1] of options A and B, conditioned on the previous choice) and the previous hidden activity 𝑍*_t_*_-1_were integrated through weight matrices 𝐔 and 𝐖 (plus bias 𝐁) to update the recurrent activity as in equation (1), where ℎ*_Z_* is a hyperbolic tangent as in equation (2). Outputs were then computed as in equation (3), where 𝐕 are output weights:

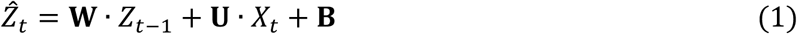

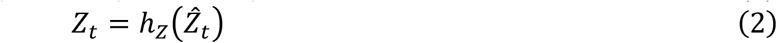

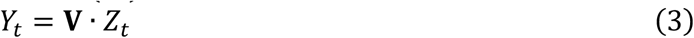

If an option was chosen, its color shade was provided as a veridical scalar value, while the unchosen option provided an ambiguous zero input. Alternatively, we also tested networks in which 𝑋*_t_* was augmented with a two-element one-hot encoding the previous choice. This alternative architecture produced results highly consistent with those reported in the main paper.

The output 𝑌*_t_* comprised three heads. The first head was passed through a sigmoid to yield action probabilities for sampling. The second and third heads were left in linear form and used only in the GUESS condition, providing state outputs (𝑠_i,t_) analogous to posterior beliefs in the cognitive model. In the MATCH task, only the first head was relevant for training. We used 64 hidden units by default and also tested smaller sizes (4, 8, 16; Supplementary Fig. S10).

### Cross-entropy loss for state prediction

In the GUESS condition, the RNN is only required to make a retrospective judgment at the end of each game: it has to predict which bag each shape was drawn from. To train the RNN, we use a cross-entropy loss function to optimize the network to match its predictions to the hidden state targets (i.e., the bag colors). Importantly, the agent needs to correctly infer the hidden states of both shapes within a game; otherwise, its performance will remain at chance level. Since the network needs to infer the hidden states of both shapes, the total loss is:

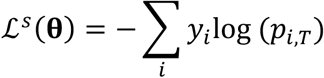

where 𝑝*_i,T_* denotes the out-of-sample *p*(state) output for option 𝑖, obtained by passing the corresponding output head through a sigmoid after the network has completed one episode of updates over trials 𝑡 = 0, …, 𝑇. 𝑦 ∈ {0, 1} is the target label for the true latent color: 0 represents the blue bag, 1 the orange bag. If the network learned to predict the correct states for both gem shapes, it would minimize this binary cross-entropy loss.

### Meta-cognitive certainty-maximization policy gradient loss

We trained the policy head to favor actions that increase a certainty signal over state representations. Let an episode have trials 𝑡 = 0, …, 𝑇. The policy 𝜋_𝛉_(𝑎*_t_*|𝑍*_T_*) outputs a Bernoulli over actions 𝑎*_t_* ∈ {0, 1} from the recurrent state 𝑍*_t_*. Here 𝛉 denotes the trainable RNN parameters. We define certainty signal as 𝜅*_t_* = min (|𝑠_1,t_|, |𝑠_2,t_|). The immediate reinforcement signal is the change in certainty that follows action 𝑎_t_, Δ𝜅 = 𝜅*_t_*_+1’_ − 𝜅*_t_*. Action on trial 𝑡 = 0 is forced to be random by design and excluded from learning. The objective maximizes expected certainty gain starting from 𝑡 = 1:

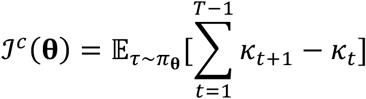

Using the policy gradient theorem with a one-step advantage, the loss we minimize is:

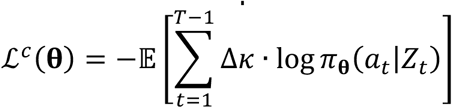

### Reward-maximization policy gradient loss

For the MATCH condition, we configured the network to make a random, unbiased choice on the first trial 𝑡 = 0. Subsequent choices were optimized with policy gradient to maximize reward accumulation. A reward 𝑟*_t_* was delivered when the network correctly pursued a pre-defined goal, such as selecting the gem shape most likely to come from the orange bag. In this formulation, the reward corresponds to the color shade outcome of the chosen option. If the target color was blue, the reward was sign-flipped relative to the color shade, ensuring alignment with the target. The policy gradient loss was then defined as:

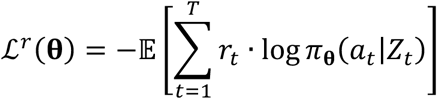

### Training procedure

The RNNs were parameterized by 𝛉 (the matrices 𝐔, 𝐖, 𝐕, and 𝐁 described in the “Neural network architecture” section) and trained using the REINFORCE algorithm^66^. This procedure optimizes the networks by differentiating the defined loss functions with respect to 𝛉: ∇_𝛉_ℒ(𝛉). In total, we trained three types of networks using different loss terms. The MATCH RNN optimized ∇_𝛉_ℒ*^T^*(𝛉); the GUESS RNN (state only) optimized ∇_𝛉_ℒ^S^(𝛉); while the GUESS meta-RNN (state + meta) optimized ∇_𝛉_[ℒ*^S^*(𝛉) + ℒ*^C^*(𝛉)].

For both conditions (MATCH and GUESS), networks were trained for 50,000 epochs. Each epoch consisted of a training batch and a test batch, both generated anew. Optimization was performed with stochastic gradient ascent using the Adam optimizer and a learning rate of 0.0001. For each network type (MATCH-orange, MATCH-blue, GUESS with state prediction only, and GUESS with state prediction plus meta-policy), 25 RNNs were trained independently from distinct small random initializations.

Training data per epoch comprised 50 sequences of variable length (2–20 trials, drawn from a truncated geometric distribution with hazard rate 0.2). Sequence lengths were sampled until all possible lengths (2 through 20) were represented at least once. For each sequence length, 200 games were simulated across the four possible color combinations for options A/B (orange/orange, blue/blue, orange/blue, blue/orange), yielding 10,000 training games per epoch. Test data followed the same structure but with fixed sequence lengths (8, 12, 16, and 20 trials) similar to the human experiment. Training with fixed sequence lengths led to results highly consistent with those reported in the main paper. For both training and test, RNNs were evaluated in mini-batches, where each mini-batch grouped games of the same sequence length. For each mini-batch, cross-entropy loss, policy gradient loss, cumulative reward, and accuracy were recorded. Asymptotic performance was reached at the end of the optimization procedure in all cases (Supplementary Fig. S9).

### Principal component analysis and effective dimensionality

Hidden-state dynamics were analyzed using Principal Component Analysis (PCA) to visualize population activity and assess dimensionality. First, independently trained RNN instances were aligned using Procrustes analysis to establish a common coordinate space by finding the optimal rotation and translation relative to a reference instance. A shared set of PCs was then derived for each network type from the concatenated, aligned hidden states, and the resulting trajectories were averaged for visualization. The intrinsic complexity was quantified using the effective dimension, or participation ratio, which measures the number of significantly active dimensions based on the squared singular values of the activity matrix, given PCA eigenvalues 𝜆*_i_*:

### Code availability

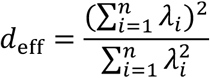

Code for computational modeling and RNNs are available at https://osf.io/q6agc/.

## Acknowledgements

This work was supported by a Starting Grant from the European Research Council (ERC-StG 759341, awarded to V.W.). Y.C. was supported by a Marie Skłodowska-Curie Fellowship (101154160). All authors benefited from an institutional grant from the Agence Nationale de la Recherche (ANR-17-EURE-0017). We thank Julien Karadayi for excellent technical support with computing cluster.

## Author contributions

Conceptualization, C.A. and V.W.; Methodology, C.A., V.W., and Y.C.; Software, J.K.L. and Y.C.; Validation, Y.C. and C.A.; Formal analysis, Y.C. and V.W.; Investigation and Data curation, Y.C., J.K.L., C.A., and I.M.; Writing – original draft, Y.C. and V.W.; Writing – review and editing, Y.C., C.A., and V.W.; Visualization, Y.C.; Supervision, V.W.; Funding acquisition, V.W.

---

**Supplementary Fig. S1.**
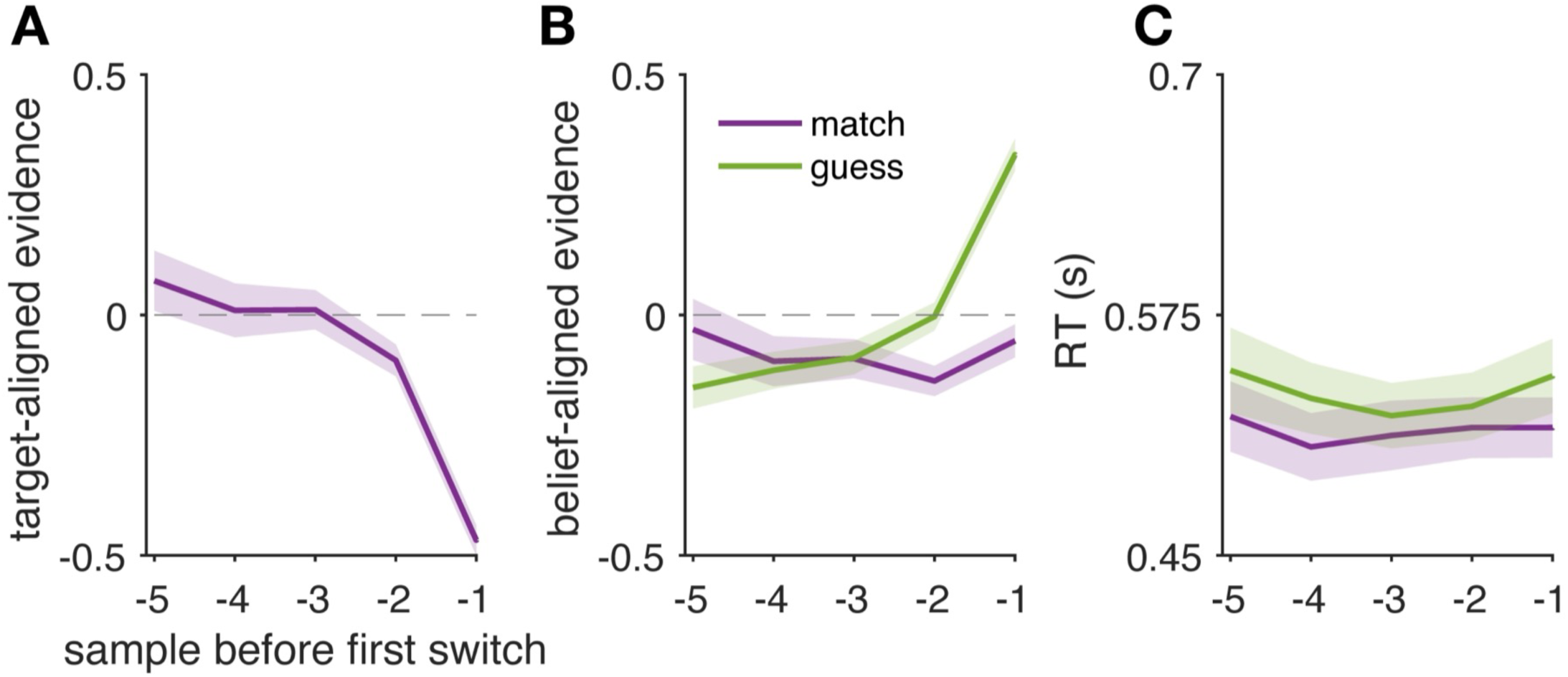
Sampling within the first early streak. Trials are time-locked to the first switch in each game. The x-axis indexes position relative to the first switch, with 0 denoting the first choice of the alternative option (not plotted) and negative indices counting backward within the initial streak. For example, the choice sequence AAAAAB is aligned so that the B trial is at 0 and the preceding A trials appear at −1, −2, and so on. Plotted are mean signed evidence (A, B) and reaction time (C) as a function of this relative index, shown separately for the two task conditions (MATCH vs. GUESS).

**Supplementary Fig. S2.**
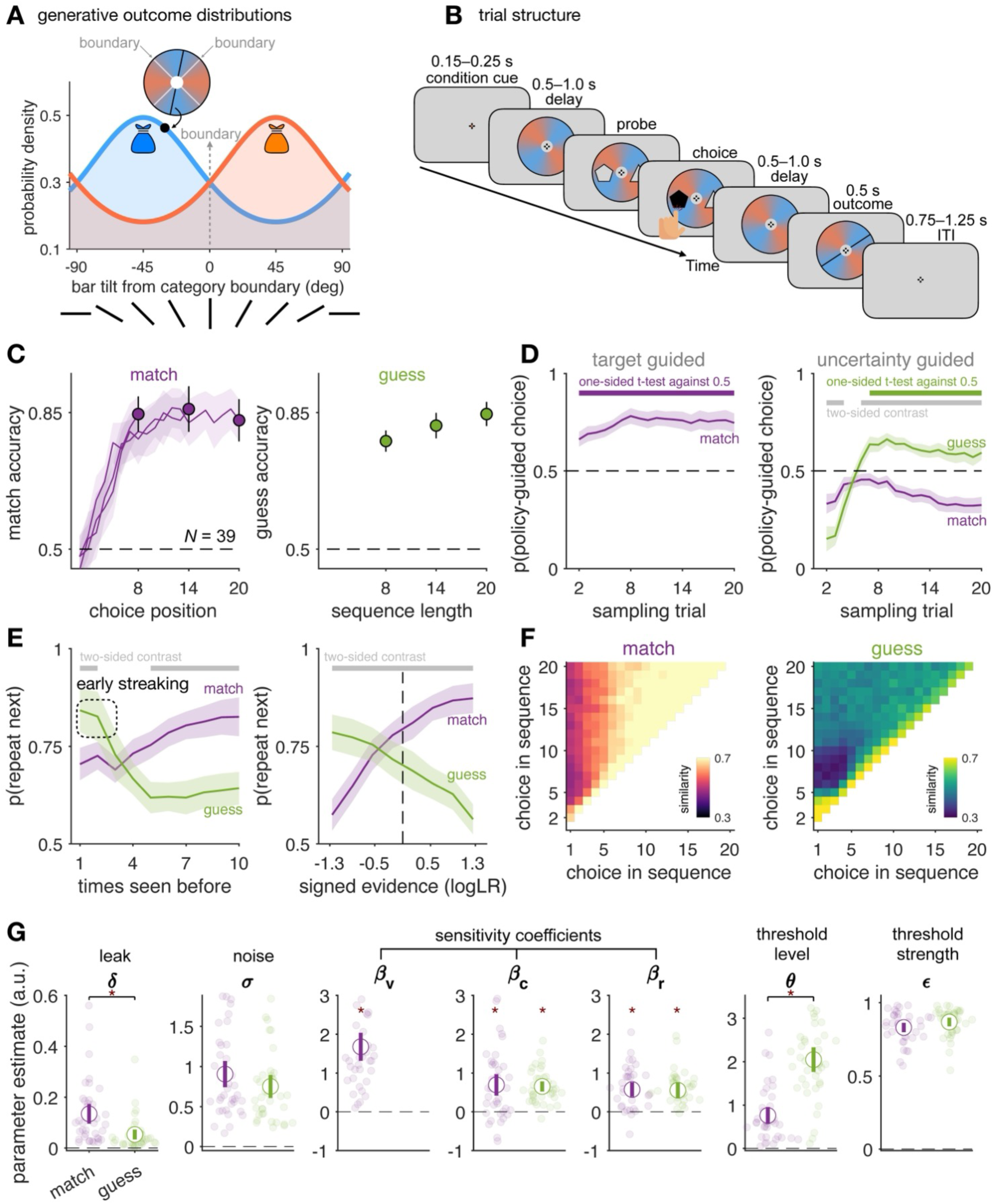
Replication study with novel stimuli. (**A**) On each trial, color outcome was drawn from a von Mises probability distribution centered either on the orientation indicated by orange or the orientation indicated by blue with a fixed concentration parameter (κ = 0.5). This setting yielded an error rate of 0.29, slightly lower than that in the main experiment (1/3). (**B**) Timing of an example trial (e.g., match orange). (**C**) Accuracy in the two conditions. (**D**) Fraction of choices aligned with directed-exploration policies across sampling trials. (**E**) Left: probability of repeating a choice as a function of how many times the current option has been sampled. Right: probability of repeating a choice as a function of signed evidence, expressed in logLR unit. Error bars show 95% CIs across N = 39 participants. (**F**) Mean choice-to-choice similarity within the sampling phase, pooled across sequences. (**G**) Estimated best-fitting model parameters. Asterisks: significant effects (max-statistic permutation tests with FWE < 0.05).

**Supplementary Fig. S3.**
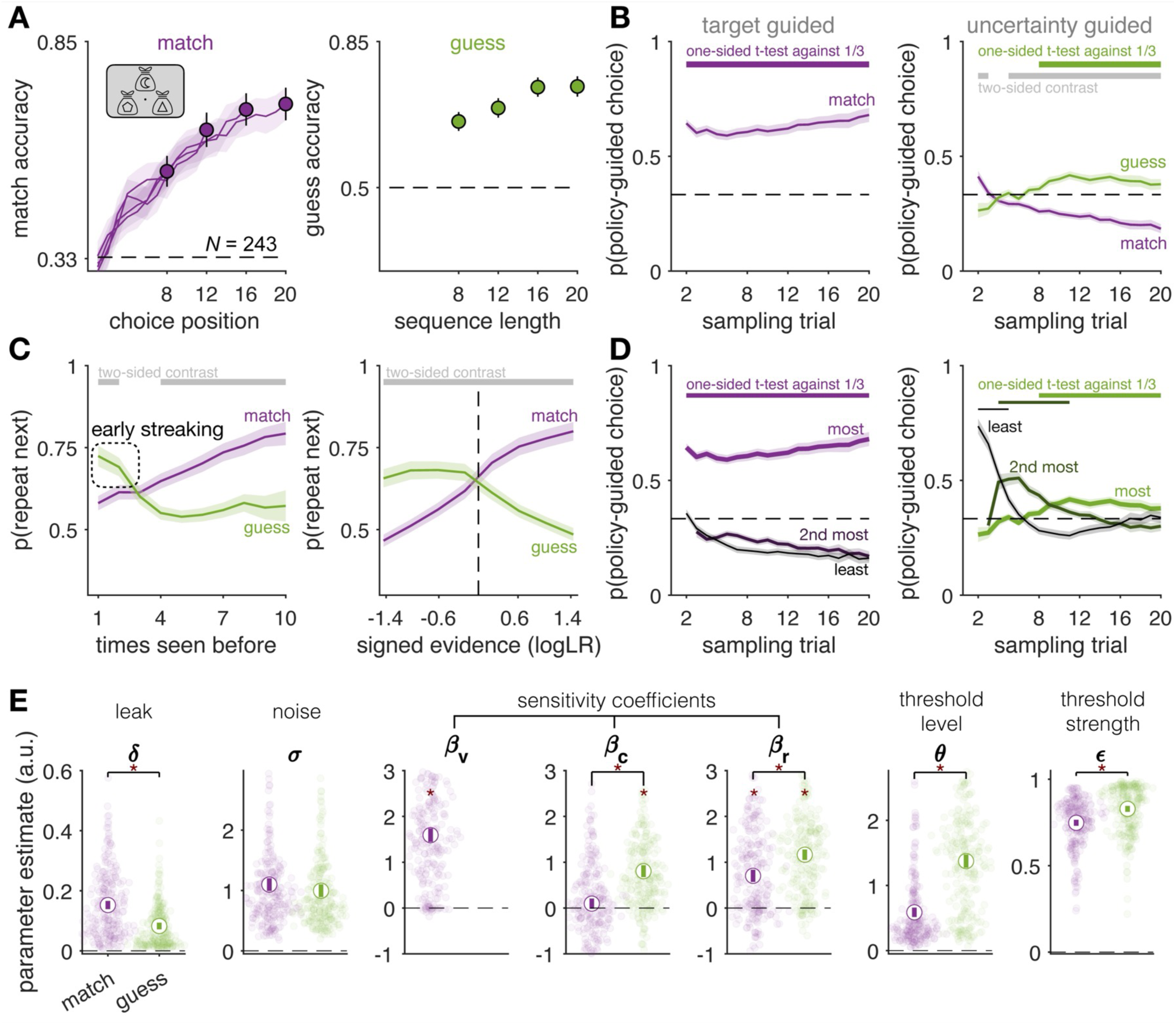
Ternary-choice study (online). Participants chose between three bags (options), each independently associated with either predominantly blue or orange shapes. Based on the smallest condition-wise contrast observed in the main study (uncertainty sensitivity *β*_c_, d_z_ = 0.251), an a priori power analysis indicated that 209 participants were required to achieve 95% power at a two-sided α = 0.05. To account for expected attrition with online datasets, we set a recruitment target of 250 participants. (**A**) Accuracy in the two conditions. In the MATCH condition, accuracy is the probability of selecting the option associated with the target color in games with counterfactual hidden structure (one bag linked to the target color, while the others to the non-target color). In the GUESS condition, accuracy is the correctness of the final probe after each game. (**B**) Fraction of choices aligned with directed-exploration policies across sampling trials. Thick lines indicate statistically significant effects (one-sided t-test for policy alignment, FDR < 0.005) and between-condition contrasts (two-sided t-test, FDR < 0.005). (**C**) Left: probability of repeating a choice as a function of how many times the current option has been sampled. Right: probability of repeating a choice as a function of signed evidence, expressed in logLR unit. In the MATCH condition, signed evidence reflects target-aligned evidence; in the GUESS condition, it reflects belief-aligned evidence, assuming an accumulation of trial-by-trial evidence for each option given the actual choices. Error bars show 95% CIs across *N* = 243 participants. (**D**) Fraction of choices consistent with the directed-exploration policy across sampling trials, shown separately for the least, second most, and most ranked options. Left panel: MATCH condition, options ranked by targetness; Right panel: GUESS condition, options ranked by uncertainty. Dashed line: chance level (1/3). (**E**) Estimated best-fitting model parameters. Asterisks indicate statistically significant contrasts between the MATCH and GUESS conditions (paired-sample t-tests, two-sided), as well as significant deviations from zero (one-sample t-tests within conditions, two-sided), both assessed using max-statistic permutation tests with family-wise error correction (FWE < 0.05).

**Supplementary Fig. S4.**
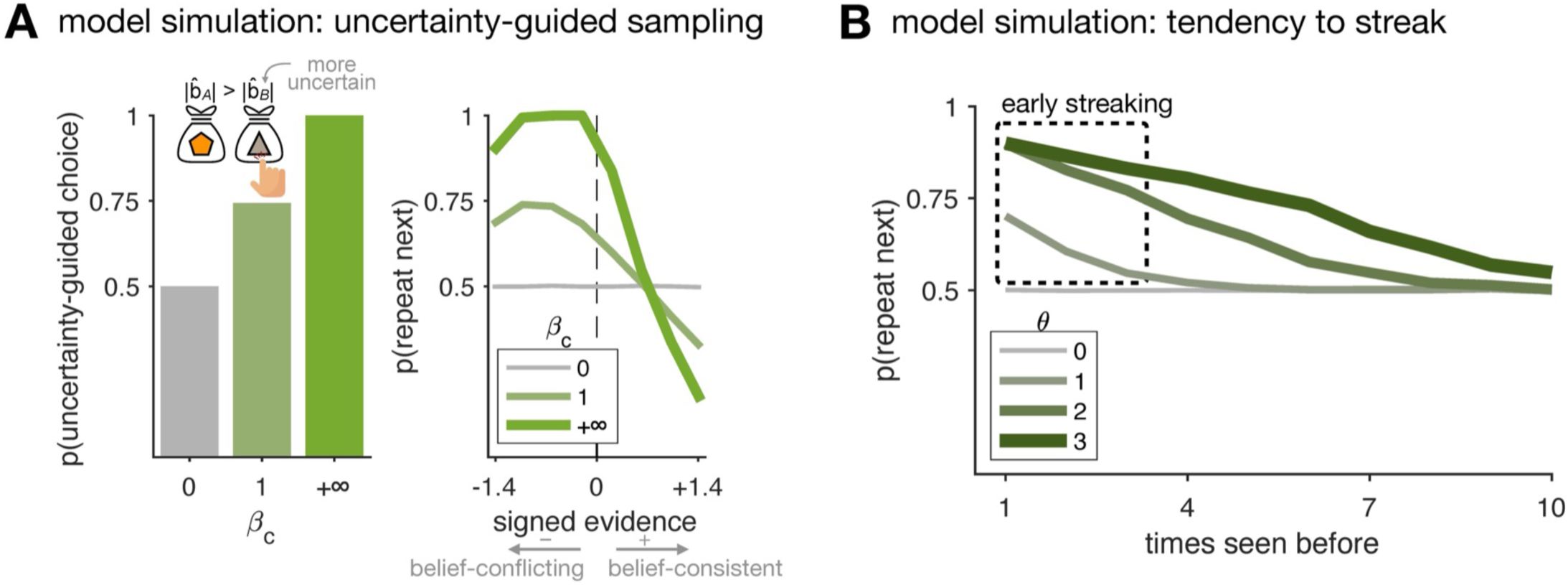
Illustrative model simulations. (A) Left: varying 𝛽_c_ changed the tendency to direct choices toward the relatively more uncertain option. 𝛽_v_, 𝛽_r_ and *θ* were set to 0; Right: a belief-aligned repetition bias emerged as a byproduct of uncertainty-guided sampling. Uncertainty is defined as the absolute cumulative logLR (belief strength) for each option. When *β*_c_ > 0 the model selects the option with the smaller belief, which reduces expected uncertainty about the bag–color mapping. A byproduct of this policy is a tendency to repeat after belief-conflicting evidence: a modestly conflicting sample pulls belief toward zero, making the just-sampled option the more uncertain one, which leads to a repeat. Belief-consistent evidence increases certainty about the sampled option, making the other option comparatively less known and prompting a switch. (B) Varying the threshold level *θ* modulated the tendency to produce early choice streaking. To isolate the threshold mechanism, the other sensitivity parameters (𝛽_v_, 𝛽_c_, and 𝛽_r_) were set to zero, repetition strength *ε* to 0.9 in these examples. Note that *θ* = 0 removes the initial sampling phase, rendering *ε* irrelevant, since repetition under the threshold never applies. The threshold *θ* explicitly controls early streaking. For *θ* > 0, the agent repeats the current option until its belief crosses *θ*, then switches and repeats the other (e.g., AAAA BBBB).

**Supplementary Fig. S5.**
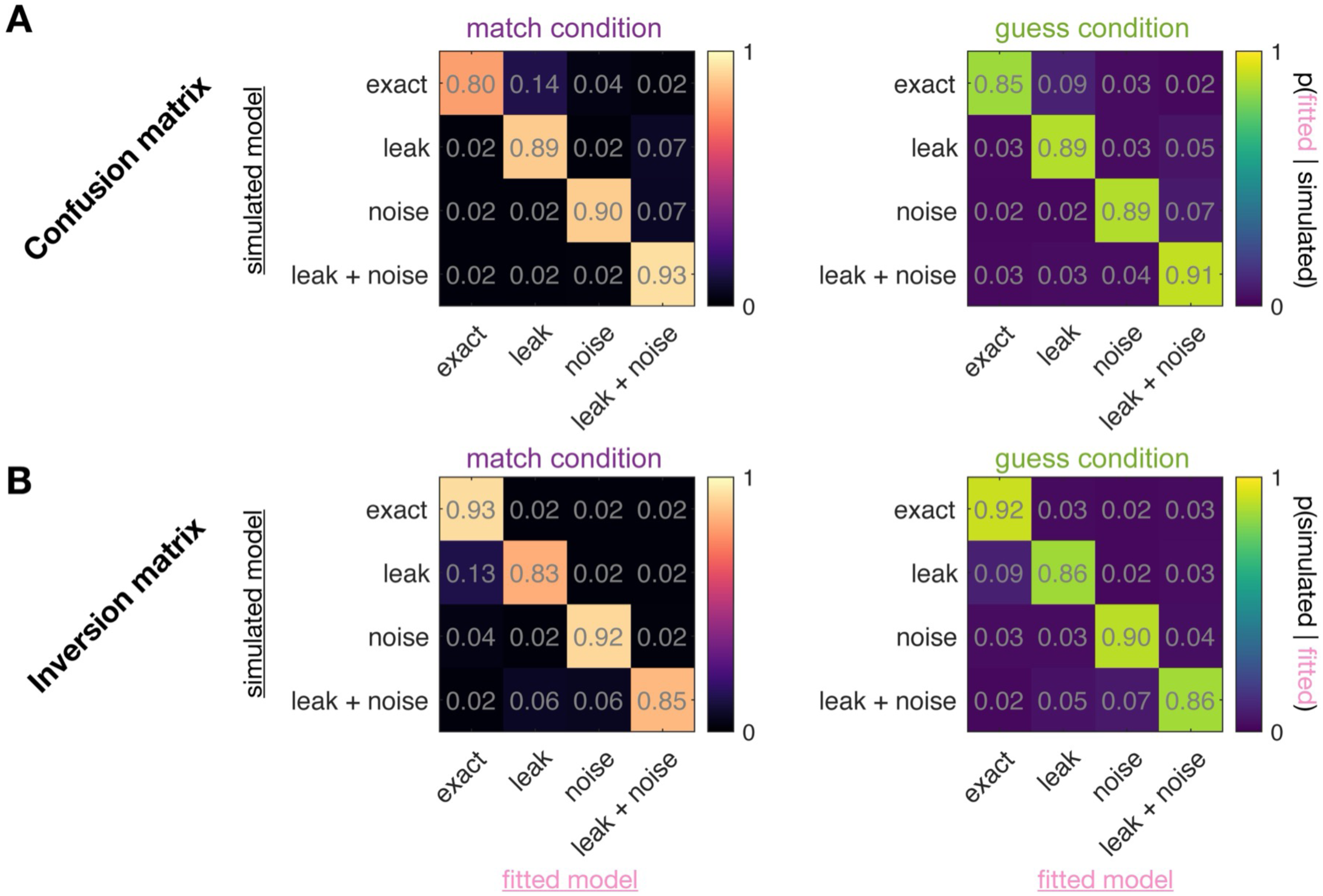
Model recovery. Using the mean best-fitting parameters from the full model across all 420 participants and both conditions (thus avoiding condition-specific bias; except with *β*_v_ = 0 in the GUESS condition), we simulated data and then fit all four model variants (full model, or knocking out leak, noise, or both) to these simulations. Color intensity indicates posterior model frequency.

**Supplementary Fig. S6.**
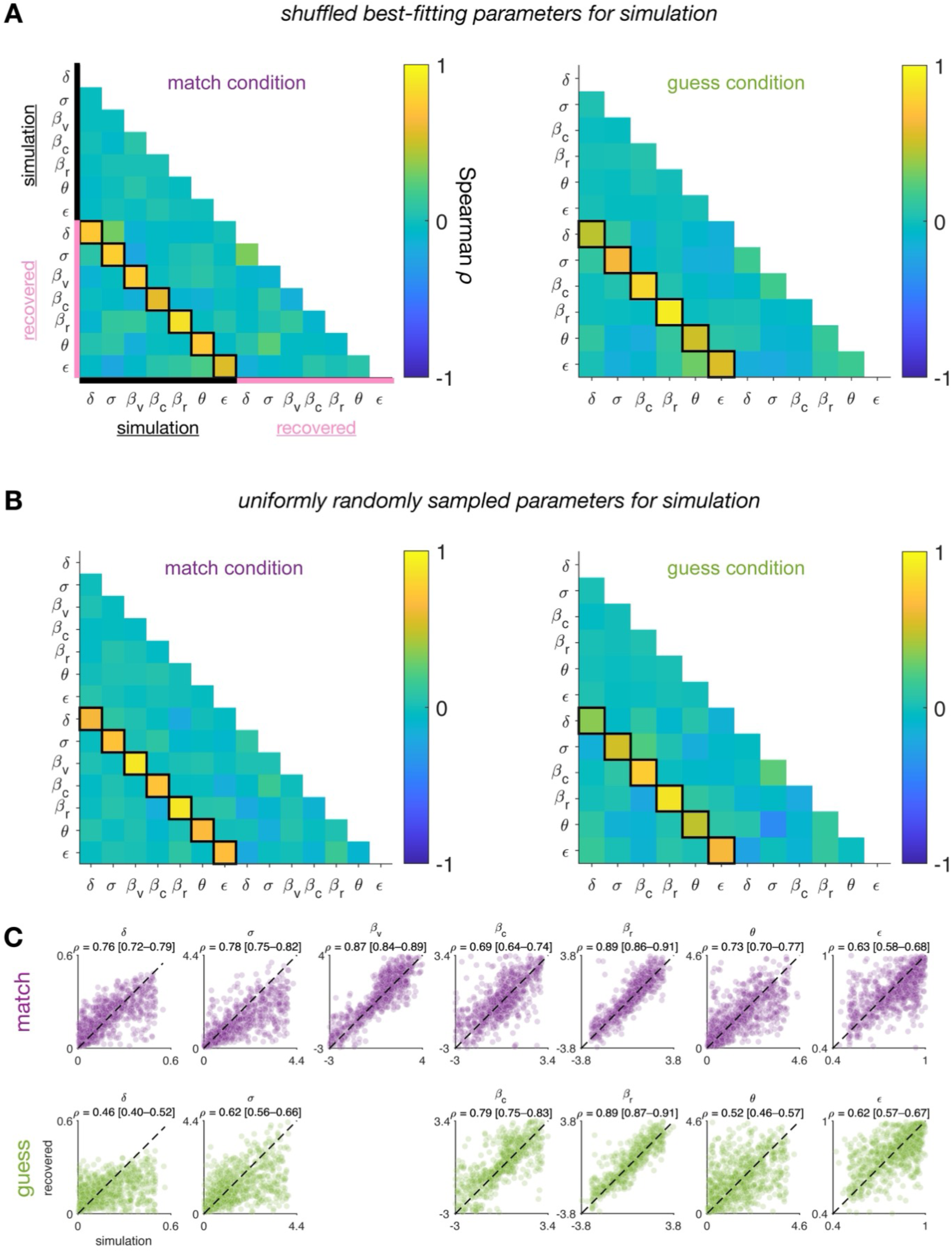
Parameter recovery. Simulated choices were generated either from participants’ best-fitting parameters (A) or from random draws within predefined ranges (*δ* ∈ [0, 0.5], *σ* ∈ [0, 4], *β*_v_, *β*_c_, *β*_r_ ∈ [−3, 3], *θ* ∈ [0, 4], *ε* ∈ [0.5, 1]; B). Correlations between generative parameters were destroyed by random permutation. Refitting the model showed no spurious correlations (A, B) and the parameters themselves were recoverable (C).

**Supplementary Fig. S7.**
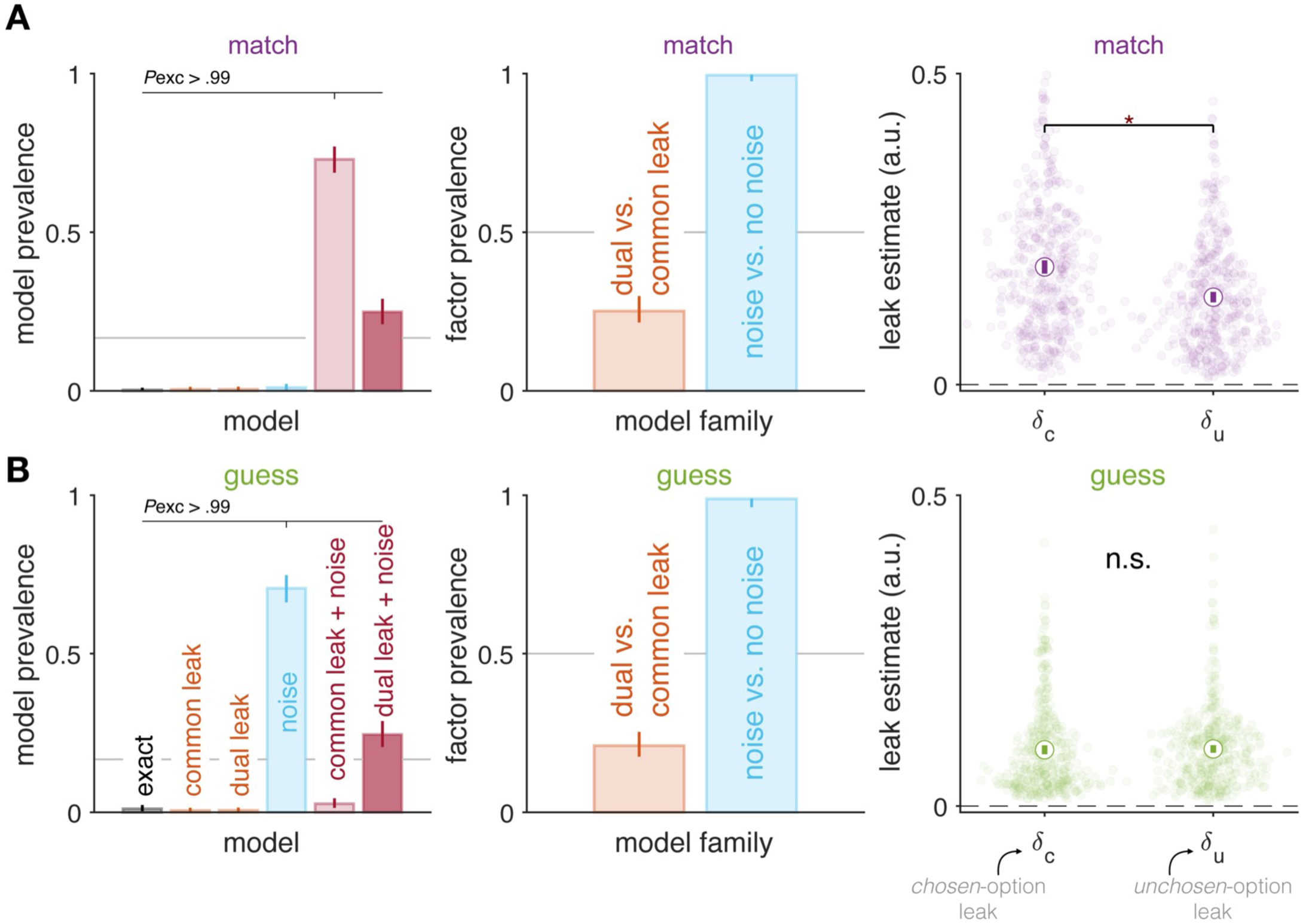
Random-effects Bayesian model comparison, including variants with independent leak parameters for the chosen and unchosen options. The model space includes single (common)-leak, dual-leak (𝛿 chosen, 𝛿 unchosen), noise-only, and full leak + noise variants. Posterior model frequencies and family-wise comparisons are computed from individual ELBOs, separately for the MATCH and GUESS conditions. Error bars indicate 95% confidence intervals.

**Supplementary Fig. S8.**
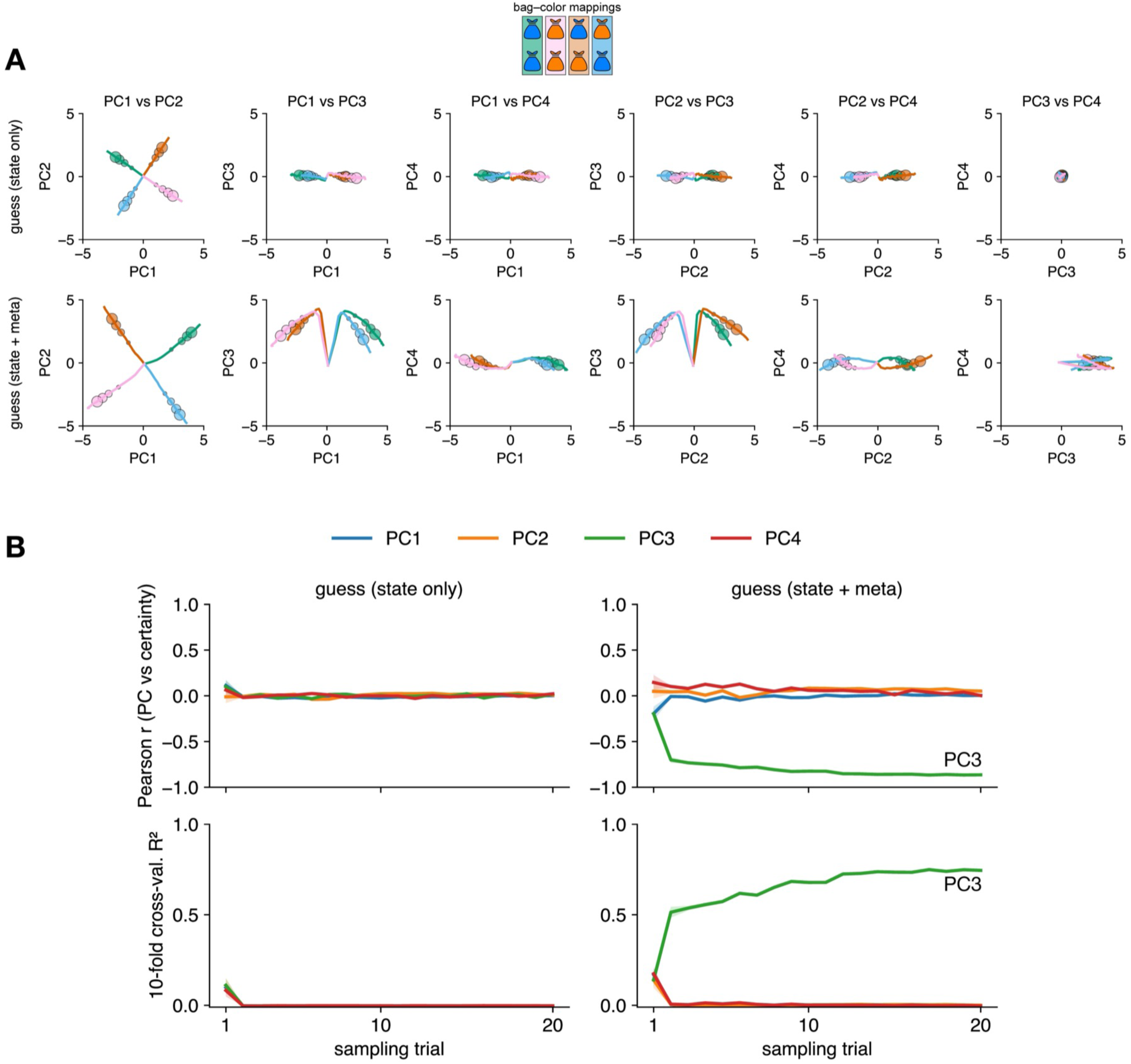
Principal Component Analysis of RNN activity. (A) Pairwise principal component plots of GUESS RNN hidden-layer activity. Curves show trajectories across sampling trials; circle markers indicate sequence lengths of 8, 12, 16, and 20. (B) Single-trial correlations (top rows) and cross-validated variance explained (bottom rows) between each PC and the network’s certainty lower bound, min(|S_A_|, |S_B_|), computed from the RNN output states.

**Supplementary Fig. S9.**
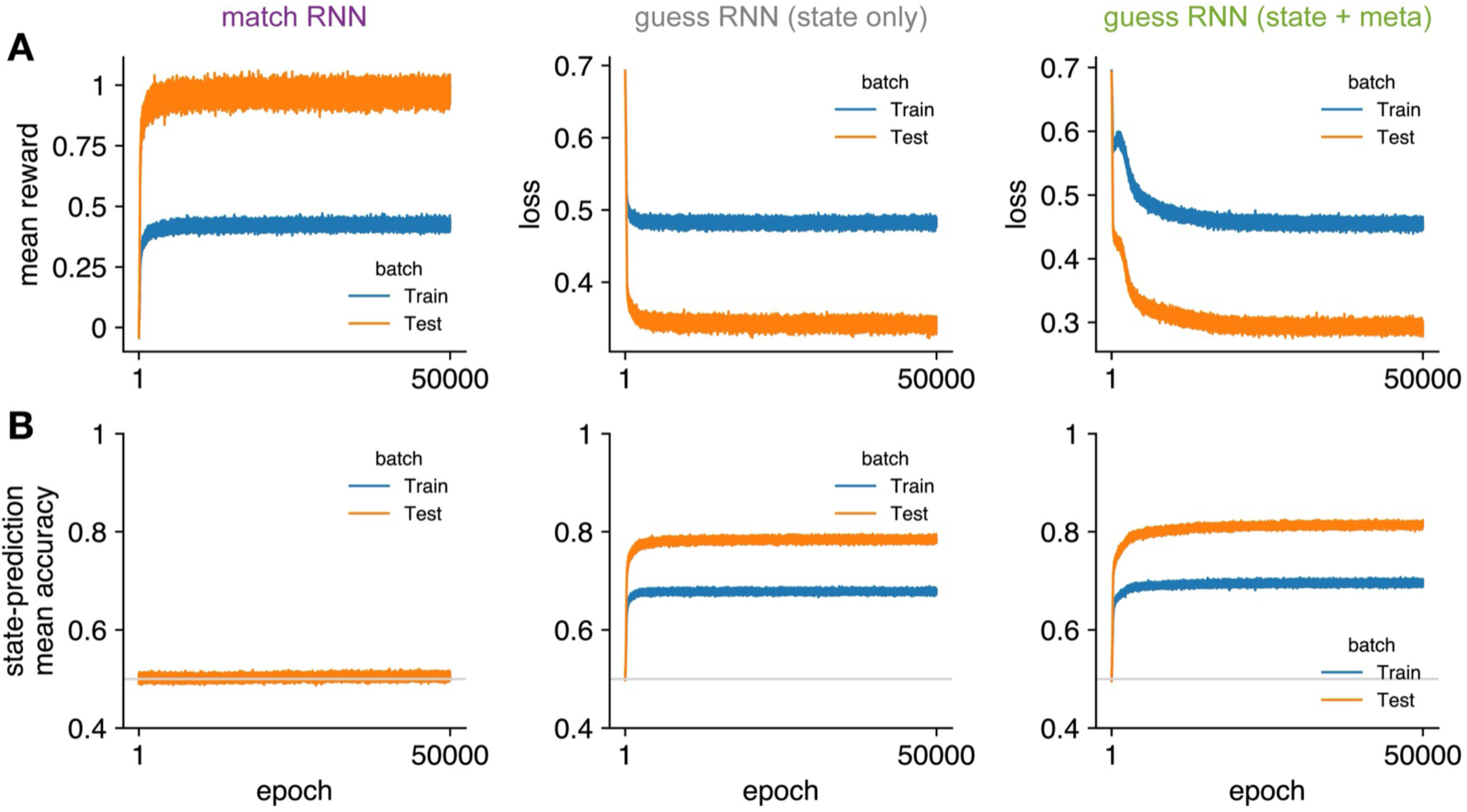
RNN convergence. (A) Training curves: mean reward per epoch for the MATCH RNN and mean loss per epoch for the GUESS RNN. (B) State-prediction accuracy as a function of training epoch, evaluated on the training and test sets. Each training epoch comprised 50 sequences with variable lengths from 2 to 20 trials, sampled from a truncated geometric distribution with hazard rate of 0.2. Sequence lengths were drawn until every length from 2 to 20 was represented at least once. Test data used fixed sequence lengths of 8, 12, 16, and 20 trials, matching the human experiment. Consequently, the training set contained, on average, a higher proportion of short sequences than the test set.

**Supplementary Fig. S10.**
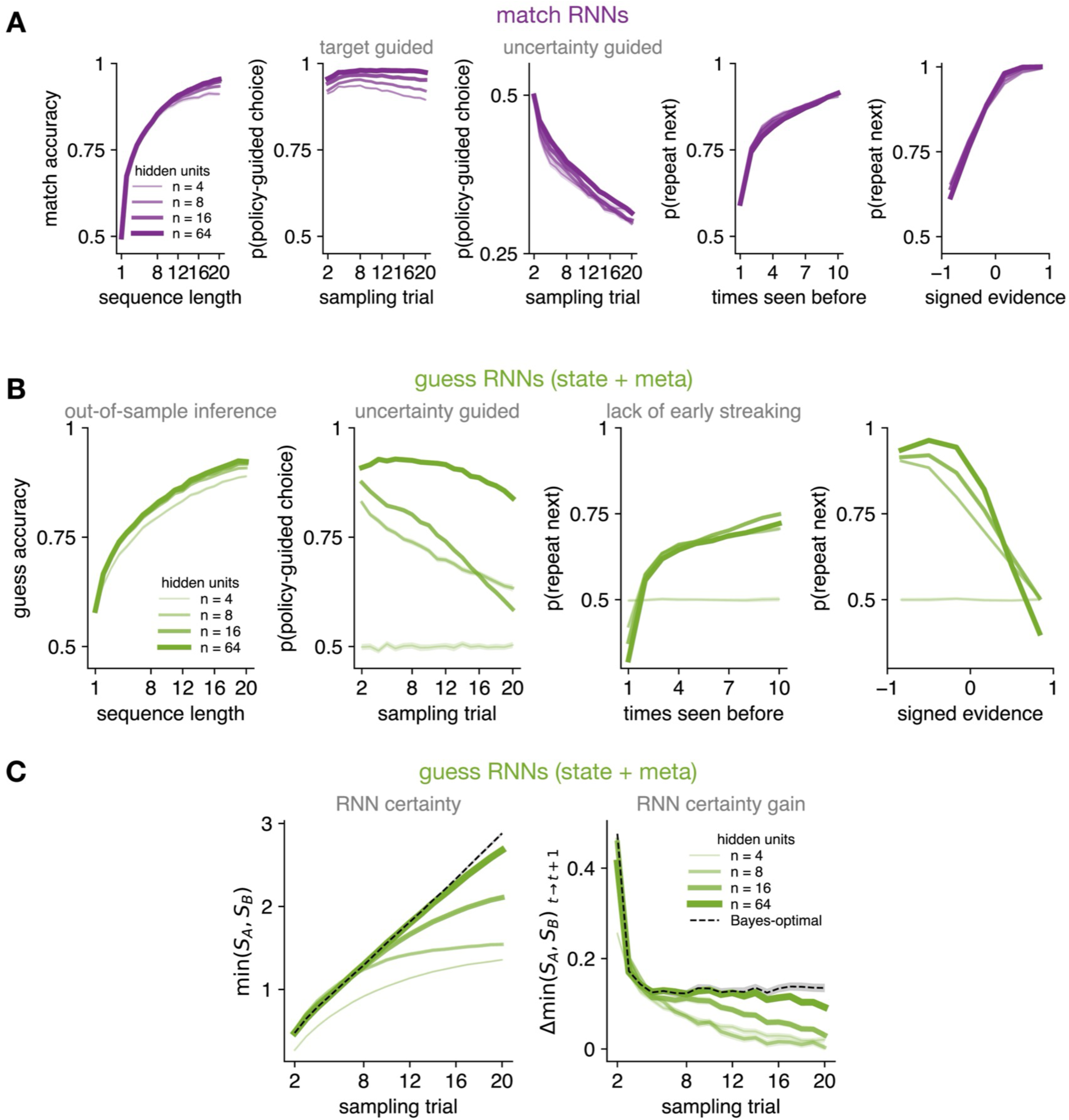
Effect of hidden-layer size on RNN performance. (A) MATCH RNNs are robust to hidden-layer size. (B) GUESS RNNs with the state plus meta-policy are sensitive to hidden-layer size for the emergence of uncertainty-guided sampling. With 4 units, the network does not exhibit uncertainty-guided sampling, whereas with as few as 8 units it does. (C) Certainty signal, i.e., network’s certainty lower bound, min(|S_A_|, |S_B_|), computed from the RNN output states. The 64-unit RNN approximates the Bayes-optimal benchmark.

**Supplementary Fig. S11.**
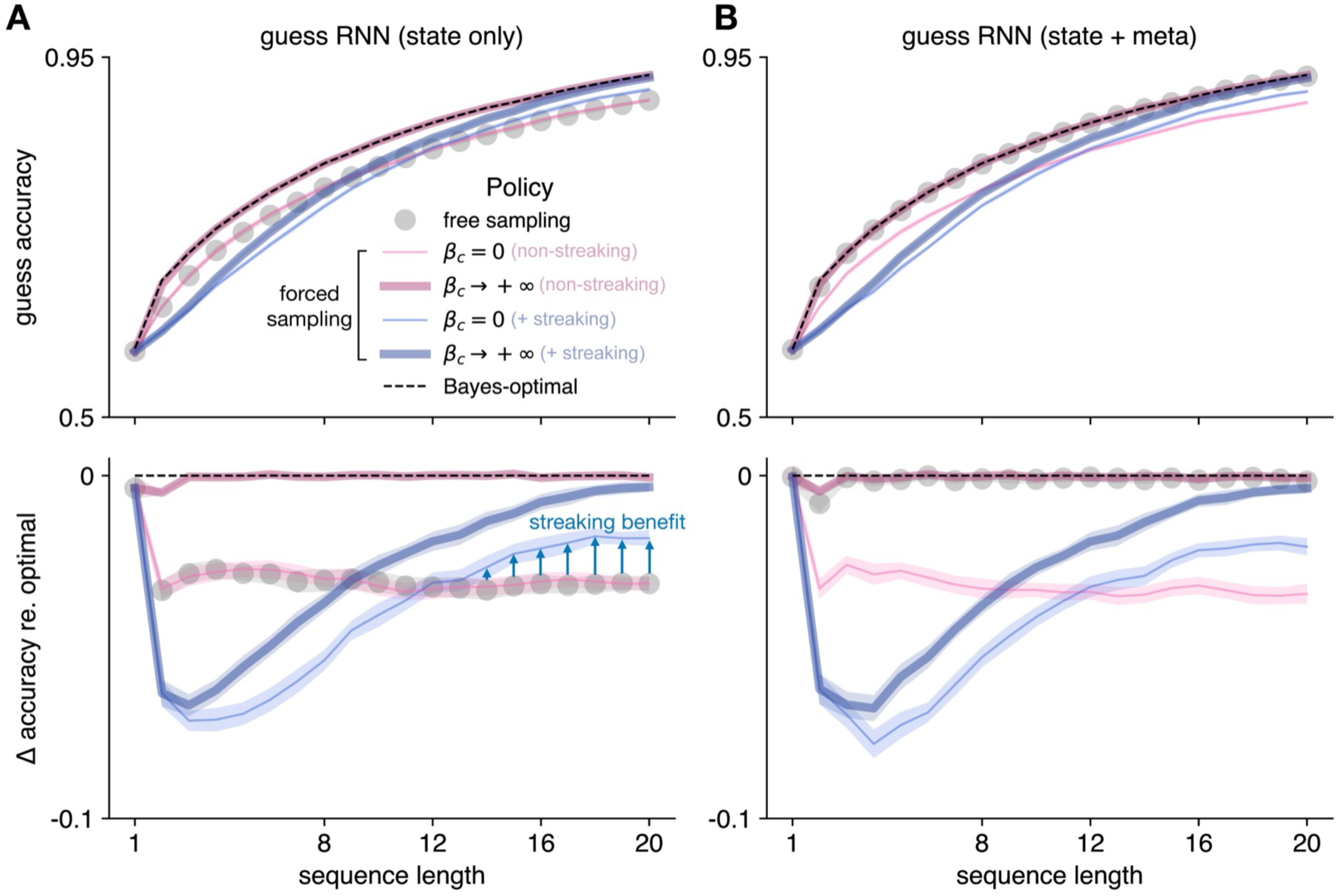
GUESS accuracy from RNNs versus a Bayes-optimal agent. (dashed). (A) GUESS RNN trained for state prediction only. (B) Meta-policy RNN trained to predict states and to increase a certainty lower bound. Free sampling: choices generated by the RNN. Forced sampling: choices follow a predefined sequence from an external agent with *θ* = 2 (streaking) or *θ* = 0 (non-streaking), and either *β*_c_ = 0 (random) or *β*_c_ → ∞ (perfect uncertainty guided). Bottom: accuracy shortfall relative to Bayes-optimal.

**Supplementary Fig. S12.**
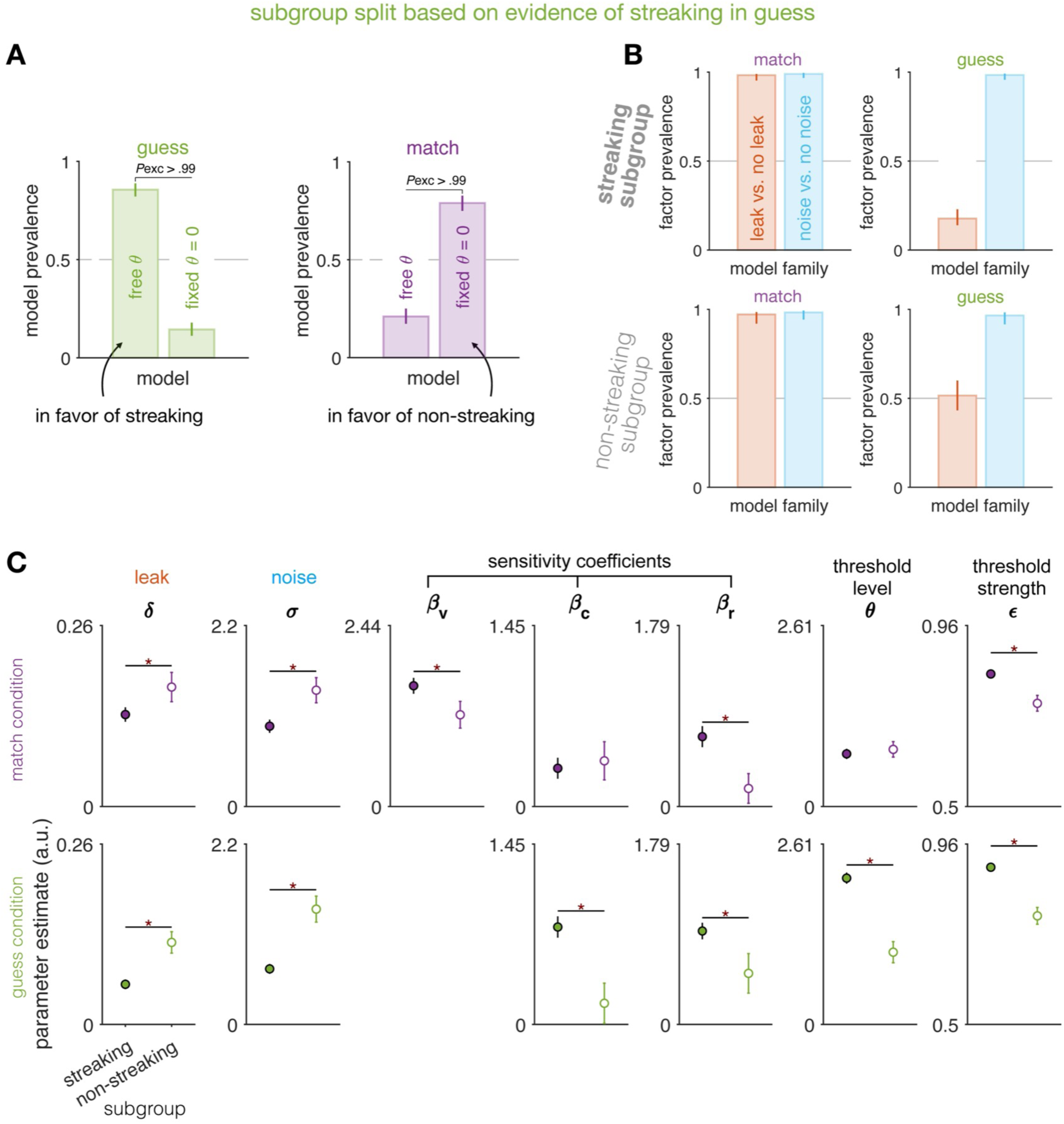
Comparison between streaking and non-streaking subgroups. (A) Random-effects model comparison between model with an initial sampling threshold and model without it, using variational ELBO as model evidence. (B) Family-wise model factor analysis within streaking subgroup and non-streaking subgroup (based on model evidence of streaking in the GUESS condition, or EoS), respectively, for each task condition (match vs. guess). (C) Parameter contrasts between the two subgroups of participants.

**Supplementary Fig. S13.**
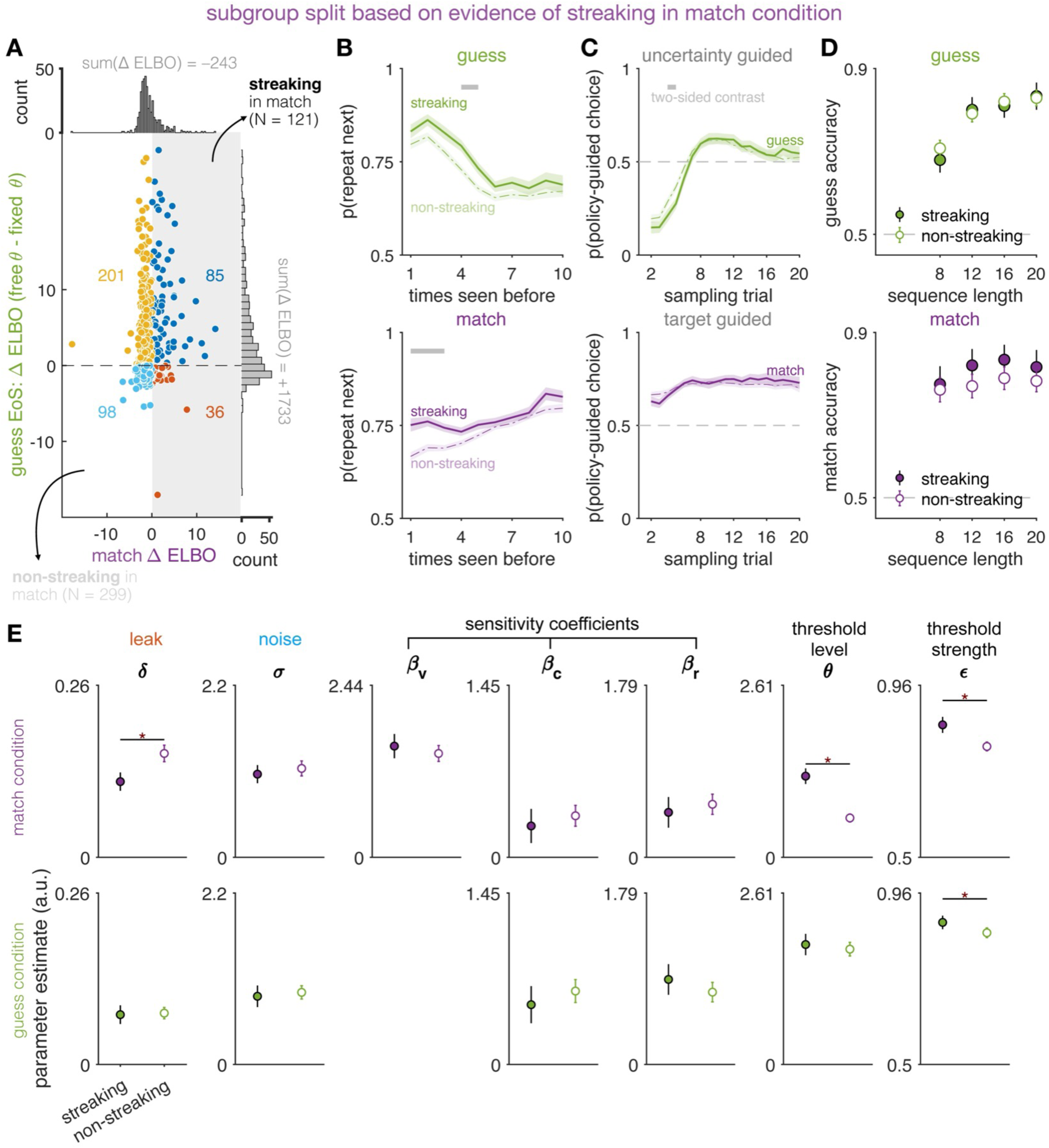
Alternative subgroup split based on model evidence for the initial phase in the MATCH condition. Conventions are otherwise the same as in Supplementary Fig. S10 and Fig. 7 of the main text.

**Supplementary Fig. S14.**
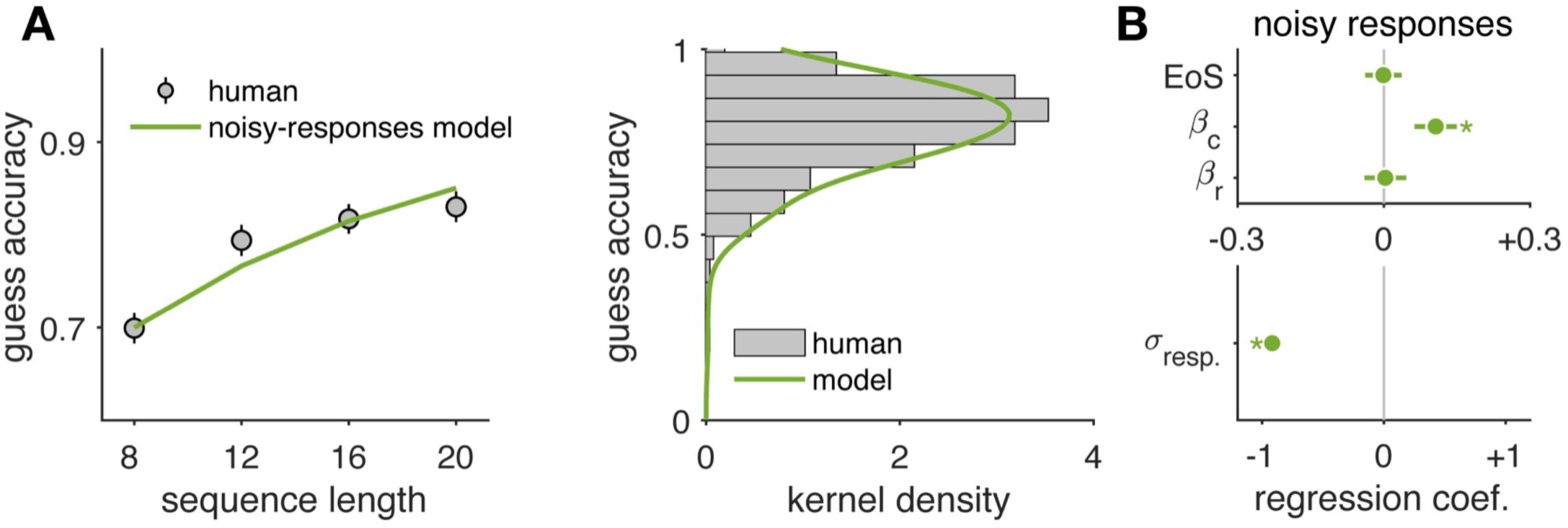
Absence of streaking benefit in a model variant with exact inferences but noisy responses. (A) GUESS accuracy from a model without learning (inference) noise; instead, action selection noise (i.e., response noise) perturbed posterior beliefs at the final probe of each game. Response-noise magnitude was drawn per participant from a gamma distribution with mean 3.5 and coefficient of variation 0.8, with parameters sampled independently to match the human distribution of mean and variance in GUESS accuracy. (B) In this model, only *β*_c_ positively predicted simulated accuracy, while EoS was not significant.

**Supplementary Fig. S15.**
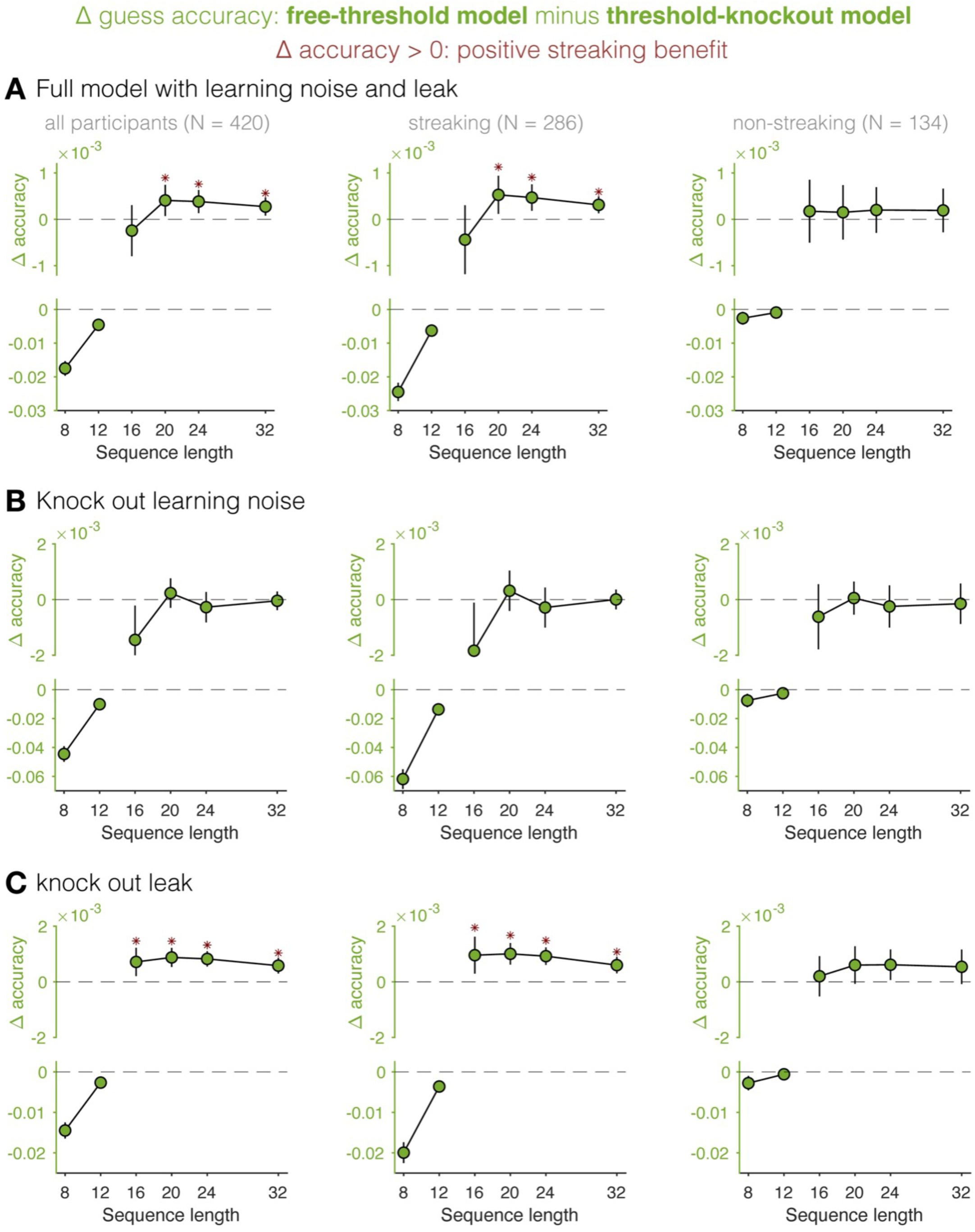
Benefit of streaking in the GUESS condition. (A) For each of 420 participants, guess accuracy was simulated (n = 1000 trajectories) using the best-fitting leaky-noisy model. A threshold-knockout model sets *θ* = 0 while keeping all other parameters identical. The streaking benefit is Δ accuracy = guess accuracy predicted by a model with threshold minus that by a model with threshold knocked out. (B) Same as A, with learning noise set to 0 in both models. (C) Same as A, with leak set to 0 in both models. For longer sequences than in the experiment, stimulus sequences were duplicated to reach lengths 24 and 32 from 12 and 16. Dark red asterisks mark one-sided t-tests against zero at p < 0.005.

**Supplementary Fig. S16.**
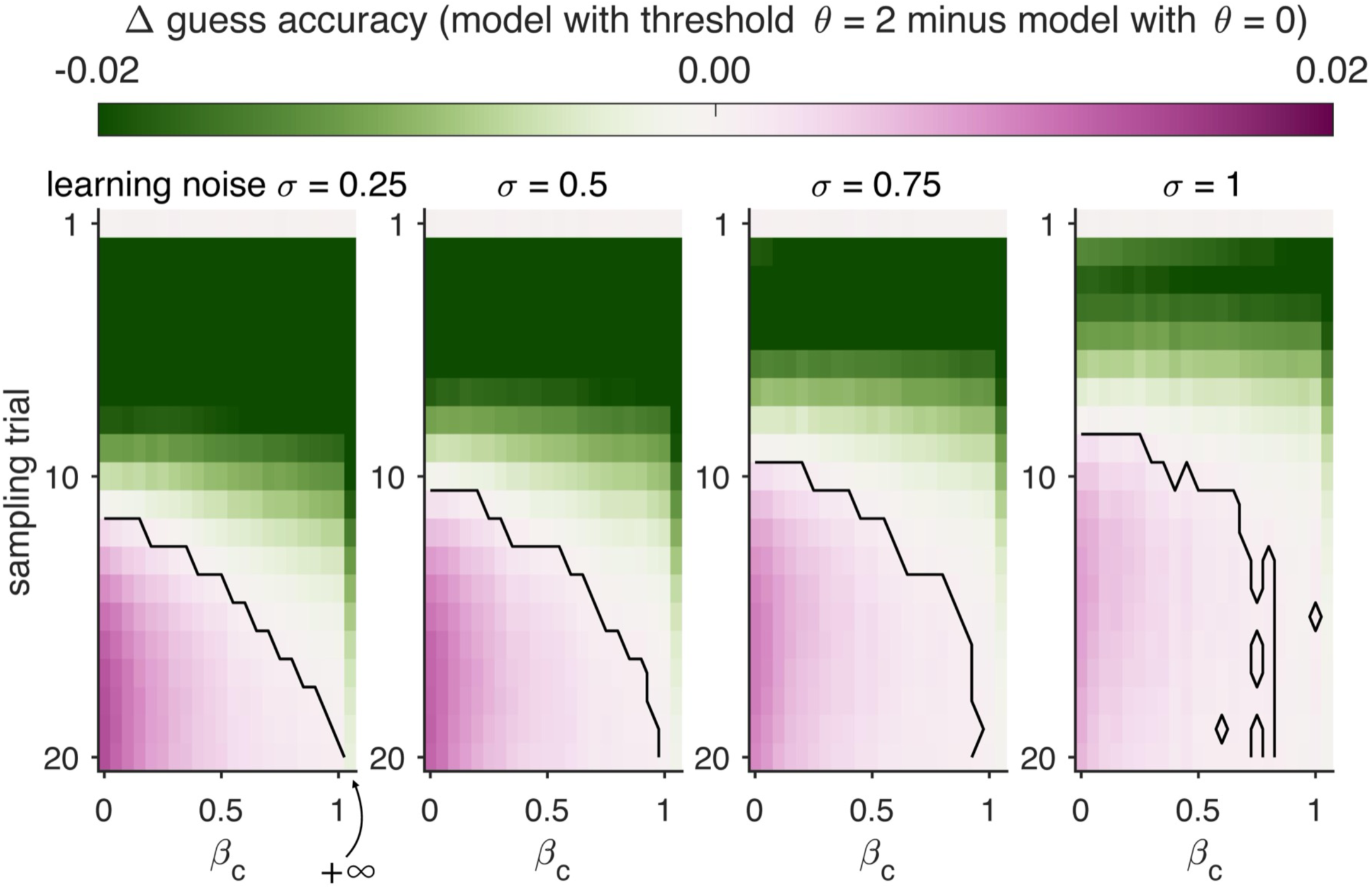
Increase in simulated accuracy from a non-streaking agent (*θ* = 0) to a streaking agent (*θ* = 2), with all other parameters fixed. Learning noise increases from left to right. The last column in each panel shows *β*_c_ → +∞, corresponding to maximal sensitivity to uncertainty-guided sampling. The solid black outline marks clusters significant by a one-sided t-test against zero at p < 0.005.

**Supplementary Fig. S17.**
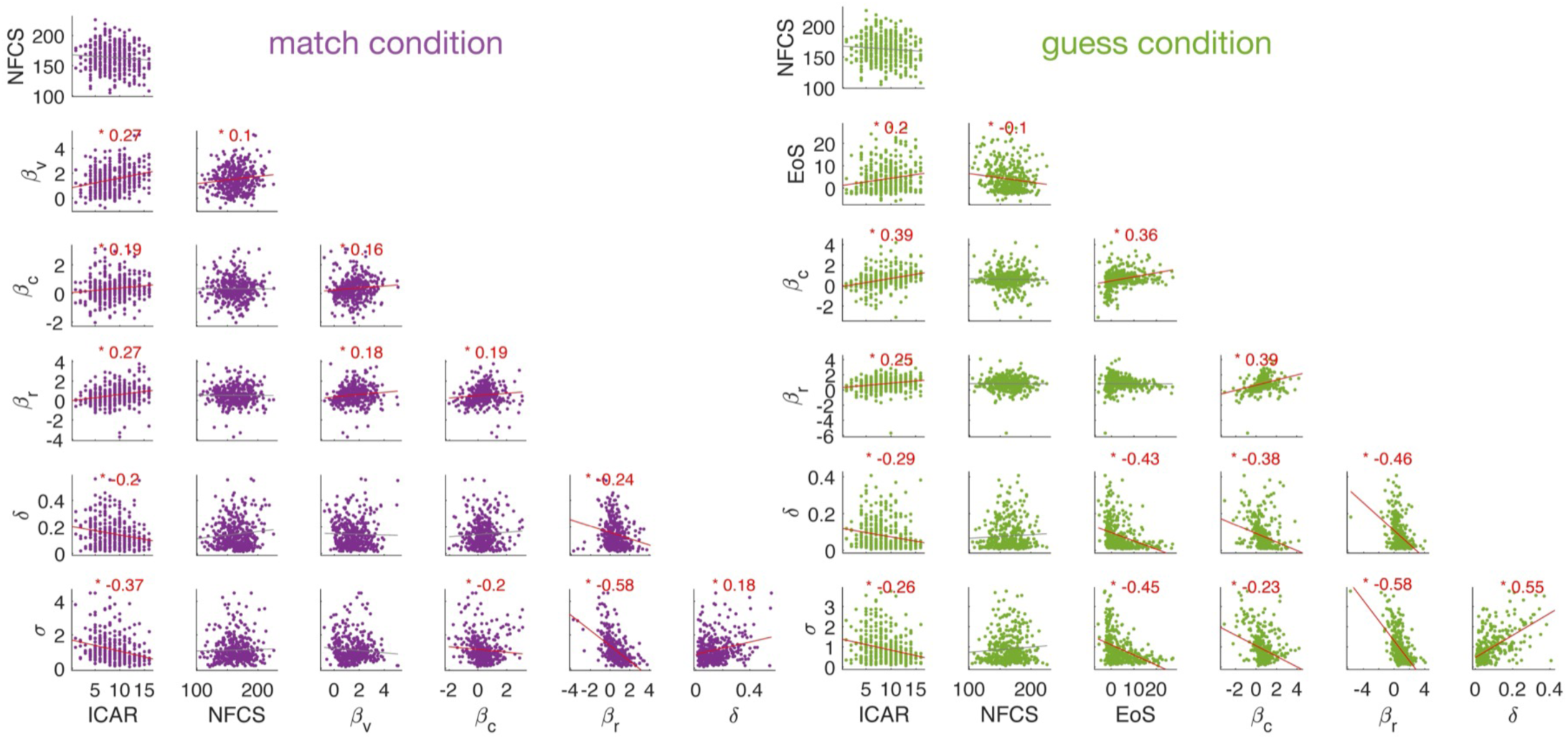
Pairwise Spearman correlations among model parameters and NFCS/ICAR. , in the MATCH and GUESS conditions, respectively.

**Supplementary Fig. S18.**
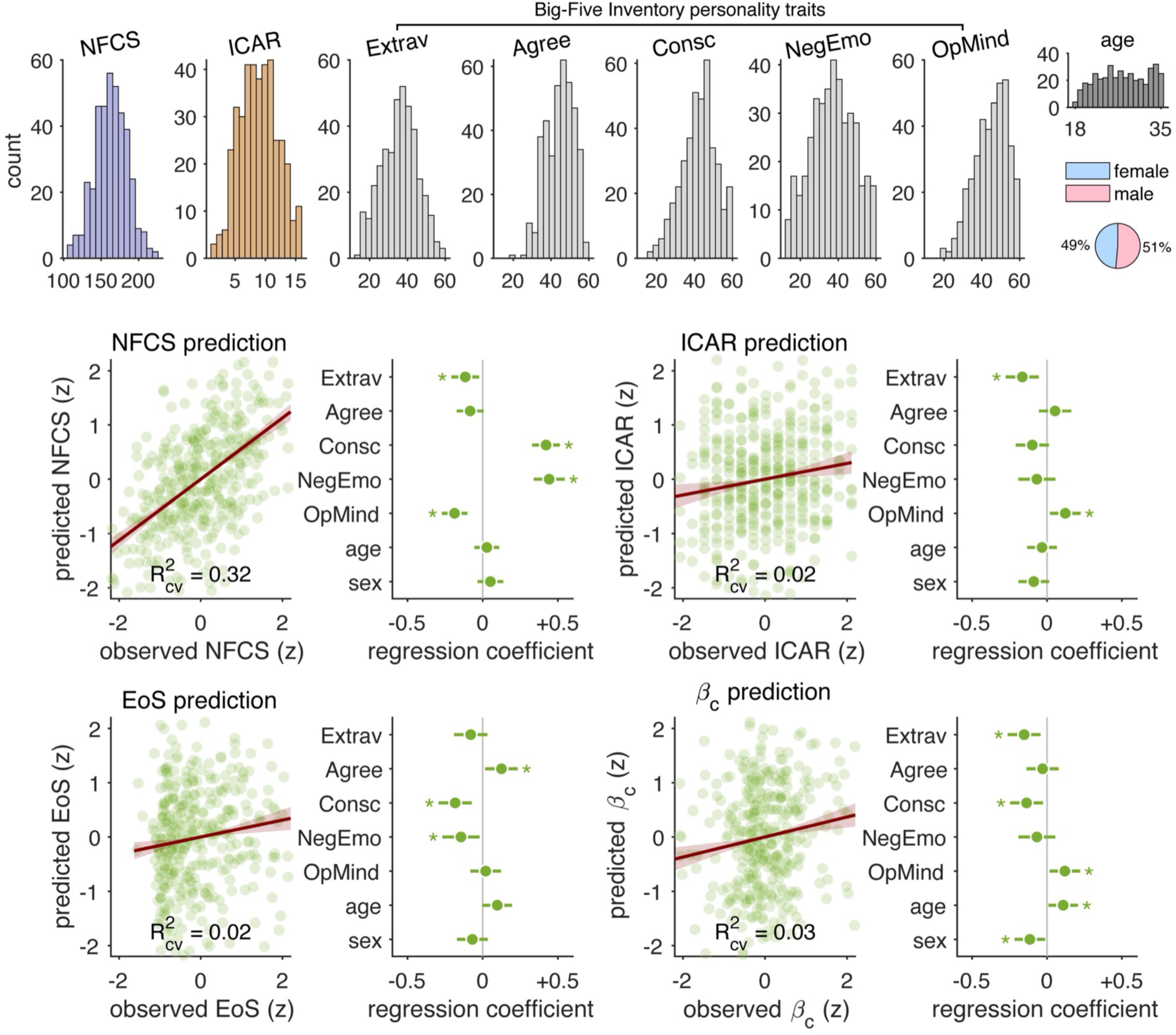
Personality trait taxonomy. Leave-one-participant-out cross-validated prediction of NFCS, ICAR, *β*_c_ (uncertainty sensitivity), and EoS (streaking tendency) from Big-Five domain scores. Extrav: Extraversion, Agree: Agreeableness, Consc: Conscientiousness, NegEmo: Negative Emotionality (formerly Neuroticism), and OpMind: Open-Mindedness (formerly Openness to Experience). Prediction panels (left, for each variable): observed values versus cross-validated predictions; the line is the bootstrap mean linear slope with a 95% CI band, and the cross-validated R² is reported at the bottom of each panel. Regression coefficient panels (right, for each variable): standardized regression coefficients by domain; the circle marks the mean coefficient, the green horizontal bar shows the 95% confidence interval, and asterisks denote coefficients different from zero (two-sided one-sample t-test).

**Supplementary Fig. S19.**
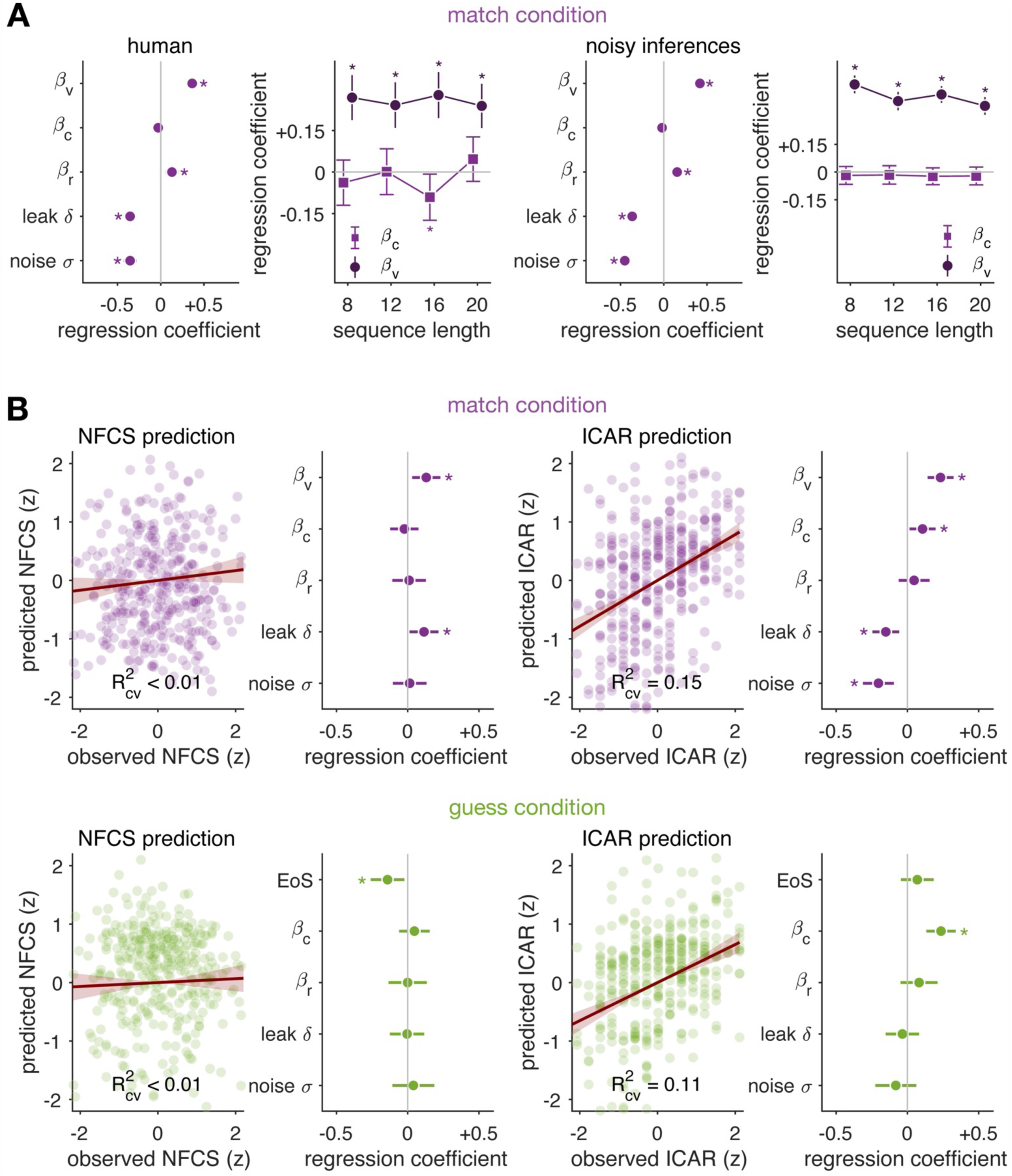
Task accuracy and psychological trait prediction from model parameters in the MATCH condition. (A) Predicting last-trial MATCH accuracy in games with one target and one non-target shape from model parameters. (B) Leave-one-participant-out cross-validated prediction of NFCS and ICAR using best-fitting parameters in the MATCH condition (top) and GUESS condition (bottom), respectively.

**Supplementary Table 1.** Human accuracy ~

| Model component | Mean coefficient | 95% CI lower | 95% CI higher | t value (df = 414) | p value |
| --- | --- | --- | --- | --- | --- |
| EoS | 0.144 | 0.046 | 0.242 | 2.896 | < 0.01* |
| $\beta_c$ | 0.211 | 0.121 | 0.300 | 4.623 | < 0.0001* |
| $\beta_r$ | 0.080 | -0.033 | 0.194 | 1.389 | 0.166 |
| leak $\delta$ | -0.132 | -0.232 | -0.032 | -2.597 | < 0.01* |
| noise $\sigma$ | -0.220 | -0.345 | -0.095 | -3.454 | < 0.001* |

Full model (noisy-inferences model) predicted accuracy ~
| Model component | Mean coefficient | 95% CI lower | 95% CI higher | t value (df = 414) | p value |
| --- | --- | --- | --- | --- | --- |
| EoS | 0.063 | 0.029 | 0.097 | 3.635 | < 0.001* |
| $\beta_c$ | 0.051 | 0.020 | 0.083 | 3.226 | < 0.01* |
| $\beta_r$ | 0.008 | -0.032 | 0.048 | 0.392 | 0.695 |
| leak $\delta$ | -0.240 | -0.275 | -0.205 | -13.533 | < 0.0001* |
| noise $\sigma$ | -0.742 | -0.786 | -0.698 | -33.451 | < 0.0001* |

Control model (noisy-responses model) predicted accuracy ~
| Model component | Mean coefficient | 95% CI lower | 95% CI higher | t value (df = 414) | p value |
| --- | --- | --- | --- | --- | --- |
| EoS | -0.001 | -0.038 | 0.036 | -0.056 | 0.956 |
| $\beta_c$ | 0.106 | 0.063 | 0.150 | 4.810 | < 0.0001* |
| $\beta_r$ | 0.003 | -0.040 | 0.046 | 0.134 | 0.893 |
| $\sigma_{resp.}$ | -0.916 | -0.953 | -0.880 | -49.637 | < 0.0001* |

NFCS ~ (N = 387; excluding 4 outliers identified by residual and influence diagnostics)
| Model component | Mean coefficient | 95% CI lower | 95% CI higher | t value (df = 381) | p value |
| --- | --- | --- | --- | --- | --- |
| EoS | -0.142 | -0.260 | -0.023 | -2.355 | 0.019* |
| $\beta_c$ | 0.047 | -0.060 | 0.154 | 0.864 | 0.388 |
| $\beta_r$ | -0.00045 | -0.134 | 0.134 | -0.007 | 0.995 |
| leak $\delta$ | -0.004 | -0.125 | 0.116 | -0.073 | 0.942 |
| noise $\sigma$ | 0.040 | -0.107 | 0.187 | 0.532 | 0.595 |

NFCS ~ (N = 391; including the four outliers)
| Model component | Mean coefficient | 95% CI lower | 95% CI higher | t value (df = 385) | p value |
| --- | --- | --- | --- | --- | --- |
| EoS | -0.126 | -0.248 | -0.004 | -2.031 | 0.042* |
| $\beta_c$ | 0.020 | -0.089 | 0.130 | 0.368 | 0.713 |
| $\beta_r$ | 0.013 | -0.125 | 0.151 | 0.190 | 0.849 |
| leak $\delta$ | -0.003 | -0.127 | 0.122 | -0.043 | 0.966 |
| noise $\sigma$ | 0.038 | -0.114 | 0.191 | 0.492 | 0.623 |

ICAR ~
| Model component | Mean coefficient | 95% CI lower | 95% CI higher | t value (df = 385) | p value |
| --- | --- | --- | --- | --- | --- |
| EoS | 0.070 | -0.045 | 0.184 | 1.191 | 0.234 |
| $\beta_c$ | 0.236 | 0.133 | 0.339 | 4.506 | < 0.0001* |
| $\beta_r$ | 0.082 | -0.048 | 0.212 | 1.241 | 0.215 |
| leak $\delta$ | -0.035 | -0.152 | 0.082 | -0.592 | 0.554 |
| noise $\sigma$ | -0.081 | -0.225 | 0.062 | -1.116 | 0.265 |

**Supplementary Table 2.** NFCS to human accuracy, mediator = EoS, control variables = {age, sex}, (N = 391)

| Path | Mean coefficient | 95% CI lower | 95% CI higher | t value | p value |
| --- | --- | --- | --- | --- | --- |
| a | -0.1294 | -0.2411 | -0.0177 | -2.28 | 0.0233* |
| b | 0.3267 | 0.2393 | 0.4141 | 7.35 | < 0.0001* |
| direct c' | -0.0046 | -0.0963 | 0.0871 | -0.10 | 0.921 |
| total | -0.0469 | -0.1441 | 0.0504 | -0.95 | 0.344 |
| indirect a*b | -0.0423 | -0.0802 | -0.0062 |  | 0.0232* |

EoS to human accuracy, mediator = NFCS, control variables = {age, sex}, (N = 391)
| Path | Mean coefficient | 95% CI lower | 95% CI higher | t value | p value |
| --- | --- | --- | --- | --- | --- |
| a | -0.1258 | -0.2354 | -0.0161 | -2.26 | 0.0247* |
| b | -0.0046 | -0.0963 | 0.0871 | -0.10 | 0.921 |
| direct c' | 0.3267 | 0.2393 | 0.4141 | 7.35 | < 0.0001* |
| total | 0.3273 | 0.2414 | 0.4132 | 7.45 | < 0.0001* |
| indirect a*b | 0.0006 | -0.0124 | 0.0139 |  | 0.913 |

ICAR to human accuracy, mediator = $\beta_c$ , control variables = {age, sex}, (N = 391)
| Path | Mean coefficient | 95% CI lower | 95% CI higher | t value | p value |
| --- | --- | --- | --- | --- | --- |
| a | 0.3135 | 0.2282 | 0.3989 | 7.22 | < 0.0001* |
| b | 0.2825 | 0.1878 | 0.3773 | 5.86 | < 0.0001* |
| direct c' | 0.3332 | 0.2398 | 0.4266 | 7.01 | < 0.0001* |
| total | 0.4217 | 0.3308 | 0.5127 | 9.12 | < 0.0001* |
| indirect a*b | 0.0886 | 0.0536 | 0.1294 |  | < 0.0001* |

$\beta_c$ to human accuracy, mediator = ICAR, control variables = {age, sex}, (N = 391)
| Path | Mean coefficient | 95% CI lower | 95% CI higher | t value | p value |
| --- | --- | --- | --- | --- | --- |
| a | 0.3155 | 0.2148 | 0.4161 | 6.16 | < 0.0001* |
| b | 0.3332 | 0.2398 | 0.4266 | 7.01 | < 0.0001* |
| direct c' | 0.2825 | 0.1878 | 0.3773 | 5.86 | < 0.0001* |
| total | 0.3876 | 0.2850 | 0.4903 | 7.43 | < 0.0001* |
| indirect a*b | 0.1051 | 0.0673 | 0.1524 |  | < 0.0001* |

**Supplementary Table 3.**

| Regressor | Mean coefficient | 95% CI lower | 95% CI higher | t value (df = 383) | p value |
| --- | --- | --- | --- | --- | --- |
| Extrav | -0.116 | -0.208 | -0.024 | -2.471 | 0.014* |
| Agree | -0.083 | -0.173 | 0.007 | -1.820 | 0.069 |
| Consc | 0.422 | 0.330 | 0.513 | 9.020 | < 0.0001* |
| NegEmo | 0.442 | 0.339 | 0.546 | 8.440 | < 0.0001* |
| OpMind | -0.187 | -0.272 | -0.102 | -4.323 | < 0.0001* |
| age | 0.028 | -0.055 | 0.110 | 0.656 | 0.512 |
| sex | 0.051 | -0.034 | 0.137 | 1.186 | 0.237 |

ICAR ~
| Regressor | Mean coefficient | 95% CI lower | 95% CI higher | t value (df = 383) | p value |
| --- | --- | --- | --- | --- | --- |
| Extrav | -0.165 | -0.276 | -0.055 | -2.936 | < 0.01* |
| Agree | 0.054 | -0.054 | 0.162 | 0.982 | 0.327 |
| Consc | -0.098 | -0.209 | 0.012 | -1.749 | 0.081 |
| NegEmo | -0.068 | -0.192 | 0.056 | -1.080 | 0.281 |
| OpMind | 0.122 | 0.020 | 0.224 | 2.347 | 0.019* |
| age | -0.033 | -0.133 | 0.066 | -0.663 | 0.508 |
| sex | -0.089 | -0.191 | 0.014 | -1.698 | 0.090 |

EoS ~
| Regressor | Mean coefficient | 95% CI lower | 95% CI higher | t value (df = 383) | p value |
| --- | --- | --- | --- | --- | --- |
| Extrav | -0.080 | -0.190 | 0.031 | -1.420 | 0.156 |
| Agree | 0.125 | 0.017 | 0.233 | 2.276 | 0.0234* |
| Consc | -0.183 | -0.293 | -0.073 | -3.259 | < 0.01* |
| NegEmo | -0.144 | -0.268 | -0.021 | -2.294 | 0.022* |
| OpMind | 0.020 | -0.082 | 0.122 | 0.384 | 0.701 |
| age | 0.097 | -0.002 | 0.196 | 1.930 | 0.054 |
| sex | -0.069 | -0.171 | 0.034 | -1.316 | 0.189 |

$\beta_c$ ~
| Regressor | Mean coefficient | 95% CI lower | 95% CI higher | t value (df = 383) | p value |
| --- | --- | --- | --- | --- | --- |
| Extrav | -0.153 | -0.263 | -0.043 | -2.731 | < 0.01* |
| Agree | -0.029 | -0.137 | 0.078 | -0.537 | 0.592 |
| Consc | -0.137 | -0.246 | -0.027 | -2.446 | 0.015* |
| NegEmo | -0.067 | -0.190 | 0.056 | -1.075 | 0.283 |
| OpMind | 0.119 | 0.018 | 0.221 | 2.308 | 0.0215* |
| age | 0.108 | 0.010 | 0.207 | 2.157 | 0.0316* |
| sex | -0.114 | -0.216 | -0.012 | -2.207 | 0.0278* |

*Big-Five domain scores:* Extrav: Extraversion, Agree: Agreeableness, Consc: Conscientiousness, NegEmo: Negative Emotionality (formerly Neuroticism), and OpMind: Open-Mindedness (formerly Openness to Experience).

## Notes

### Competing Interest Statement

The authors have declared no competing interest.

### Summary of Updates

We combine our original dataset (N = 420) with two additional experiments (total N = 282) that corroborate the behavioral signatures of the 'streak-first, explore-second' dual policy and support its robustness and generalization. We describe the results of these additional experiments in two supplementary figures (Supplementary Fig. S2 and S3).

## References

1. Gittins, J. C. Bandit processes and dynamic allocation indices. J. R. Stat. Soc. Series B Stat. Methodol. 41, 148–164 (1979).

2. Mehlhorn, K. et al. Unpacking the exploration–exploitation tradeoff: A synthesis of human and animal literatures. Decision (Wash., DC) 2, 191–215 (2015).

3. Sutton, R. S. & Barto, A. G. Reinforcement learning: an introduction MIT Press. Cambridge, MA (1998).

4. Daw, N. D., O’Doherty, J. P., Dayan, P., Seymour, B. & Dolan, R. J. Cortical substrates for exploratory decisions in humans. Nature 441, 876–879 (2006).

5. Cohen, J. D., McClure, S. M. & Yu, A. J. Should I stay or should I go? How the human brain manages the trade-off between exploitation and exploration. Philos. Trans. R. Soc. Lond. B Biol. Sci. 362, 933–942 (2007).

6. Schulz, E., Franklin, N. T. & Gershman, S. J. Finding structure in multi-armed bandits. Cogn. Psychol. 119, 101261 (2020).

7. Wulff, D. U., Mergenthaler-Canseco, M. & Hertwig, R. A meta-analytic review of two modes of learning and the description-experience gap. Psychol. Bull. 144, 140–176 (2018).

8. Wulff, D. U., Hills, T. T. & Hertwig, R. How short- and long-run aspirations impact search and choice in decisions from experience. Cognition 144, 29–37 (2015).

9. Wilson, R. C., Bonawitz, E., Costa, V. D. & Ebitz, R. B. Balancing exploration and exploitation with information and randomization. Curr Opin Behav Sci 38, 49–56 (2021).

10. Hills, T. T. & Hertwig, R. Information search in decisions from experience. Do our patterns of sampling foreshadow our decisions?: Do our patterns of sampling foreshadow our decisions? Psychol. Sci. 21, 1787–1792 (2010).

11. Wilson, R. C., Geana, A., White, J. M., Ludvig, E. A. & Cohen, J. D. Humans use directed and random exploration to solve the explore-exploit dilemma. J. Exp. Psychol. Gen. 143, 2074–2081 (2014).

12. Sharot, T. & Sunstein, C. R. How people decide what they want to know. *Nat*. Hum. Behav. 4, 14–19 (2020).

13. Bromberg-Martin, E. S. & Hikosaka, O. Midbrain dopamine neurons signal preference for advance information about upcoming rewards. Neuron 63, 119–126 (2009).

14. Bromberg-Martin, E. S. et al. A neural mechanism for conserved value computations integrating information and rewards. Nat. Neurosci. 27, 159–175 (2024).

15. Blanchard, T. C., Hayden, B. Y. & Bromberg-Martin, E. S. Orbitofrontal cortex uses distinct codes for different choice attributes in decisions motivated by curiosity. Neuron 85, 602–614 (2015).

16. Kang, M. J. et al. The wick in the candle of learning: epistemic curiosity activates reward circuitry and enhances memory: Epistemic curiosity activates reward circuitry and enhances memory. Psychol. Sci. 20, 963–973 (2009).

17. Charpentier, C. J., Bromberg-Martin, E. S. & Sharot, T. Valuation of knowledge and ignorance in mesolimbic reward circuitry. Proc. Natl. Acad. Sci. U. S. A. 115, E7255–E7264 (2018).

18. Kobayashi, K. & Kable, J. W. Neural mechanisms of information seeking. Neuron (2024) doi:10.1016/j.neuron.2024.04.008.

19. Monosov, I. E. Curiosity: primate neural circuits for novelty and information seeking. Nat. Rev. Neurosci. (2024) doi:10.1038/s41583-023-00784-9.

20. Gopnik, A. Childhood as a solution to explore-exploit tensions. Philos. Trans. R. Soc. Lond. B Biol. Sci. 375, 20190502 (2020).

21. Alméras, C., Chambon, V. & Wyart, V. Competing cognitive pressures on human exploration in the absence of trade-off with exploitation. PsyArXiv (2022) doi:10.31234/osf.io/9qpuz.

22. Urai, A. E., de Gee, J. W., Tsetsos, K. & Donner, T. H. Choice history biases subsequent evidence accumulation. Elife 8, e46331 (2019).

23. Drugowitsch, J., Wyart, V., Devauchelle, A.-D. & Koechlin, E. Computational Precision of Mental Inference as Critical Source of Human Choice Suboptimality. Neuron 92, 1398–1411 (2016).

24. Lee, J. K., Rouault, M. & Wyart, V. Adaptive tuning of human learning and choice variability to unexpected uncertainty. Sci Adv 9, eadd0501 (2023).

25. Palminteri, S., Wyart, V. & Koechlin, E. The Importance of Falsification in Computational Cognitive Modeling. Trends Cogn. Sci. 21, 425–433 (2017).

26. Wilson, R. C. & Collins, A. G. E. Ten simple rules for the computational modeling of behavioral data. Elife 8, e49547 (2019).

27. Acerbi, L. Variational Bayesian Monte Carlo with noisy likelihoods. Neural Inf Process Syst abs/2006.08655, 8211–8222 (2020).

28. Collins, J., Sohl-Dickstein, J. & Sussillo, D. Capacity and trainability in recurrent neural networks. arXiv [stat.ML] (2016).

29. Acerbi, L. Variational Bayesian Monte Carlo. arXiv [stat.ML] (2018).

30. Bainbridge, T. F., Ludeke, S. G. & Smillie, L. D. Evaluating the Big Five as an organizing framework for commonly used psychological trait scales. J. Pers. Soc. Psychol. 122, 749–777 (2022).

31. Soto, C. J. & John, O. P. The next Big Five Inventory (BFI-2): Developing and assessing a hierarchical model with 15 facets to enhance bandwidth, fidelity, and predictive power. J. Pers. Soc. Psychol. 113, 117–143 (2017).

32. Gottlieb, J. & Oudeyer, P.-Y. Towards a neuroscience of active sampling and curiosity. Nat. Rev. Neurosci. 19, 758–770 (2018).

33. Bromberg-Martin, E. S., Matsumoto, M. & Hikosaka, O. Dopamine in motivational control: rewarding, aversive, and alerting. Neuron 68, 815–834 (2010).

34. Daddaoua, N., Lopes, M. & Gottlieb, J. Intrinsically motivated oculomotor exploration guided by uncertainty reduction and conditioned reinforcement in non-human primates. Sci. Rep. 6, 20202 (2016).

35. FitzGerald, T. H. B., Dolan, R. J. & Friston, K. Dopamine, reward learning, and active inference. Front. Comput. Neurosci. 9, 136 (2015).

36. Kidd, C. & Hayden, B. Y. The psychology and neuroscience of curiosity. Neuron 88, 449–460 (2015).

37. Ten, A., Kaushik, P., Oudeyer, P.-Y. & Gottlieb, J. Humans monitor learning progress in curiosity-driven exploration. Nat. Commun. 12, 5972 (2021).

38. Monsell, S. Task switching. Trends Cogn. Sci. 7, 134–140 (2003).

39. Kool, W., McGuire, J. T., Rosen, Z. B. & Botvinick, M. M. Decision making and the avoidance of cognitive demand. J. Exp. Psychol. Gen. 139, 665–682 (2010).

40. Lieder, F. & Griffiths, T. L. Resource-rational analysis: Understanding human cognition as the optimal use of limited computational resources. Behav. Brain Sci. 43, e1 (2019).

41. Markant, D. B. & Gureckis, T. M. Is it better to select or to receive? Learning via active and passive hypothesis testing. J. Exp. Psychol. Gen. 143, 94–122 (2014).

42. Findling, C., Skvortsova, V., Dromnelle, R., Palminteri, S. & Wyart, V. Computational noise in reward-guided learning drives behavioral variability in volatile environments. Nat. Neurosci. 22, 2066–2077 (2019).

43. Drevet, J., Drugowitsch, J. & Wyart, V. Efficient stabilization of imprecise statistical inference through conditional belief updating. *Nat*. Hum. Behav. 6, 1691–1704 (2022).

44. Tsetsos, K. et al. Economic irrationality is optimal during noisy decision making. Proc. Natl. Acad. Sci. U. S. A. 113, 3102–3107 (2016).

45. Findling, C. & Wyart, V. Computation noise in human learning and decision-making: origin, impact, function. Current Opinion in Behavioral Sciences 38, 124–132 (2021).

46. Weiss, A., Chambon, V., Lee, J. K., Drugowitsch, J. & Wyart, V. Interacting with volatile environments stabilizes hidden-state inference and its brain signatures. Nat. Commun. 12, 2228 (2021).

47. Beukers, A. O. et al. Blocked training facilitates learning of multiple schemas. Commun. Psychol. 2, 28 (2024).

48. Dekker, R. B., Otto, F. & Summerfield, C. Curriculum learning for human compositional generalization. Proc. Natl. Acad. Sci. U. S. A. 119, e2205582119 (2022).

49. Collins, A. G. E. & Frank, M. J. Within- and across-trial dynamics of human EEG reveal cooperative interplay between reinforcement learning and working memory. Proc. Natl. Acad. Sci. U. S. A. 115, 2502–2507 (2018).

50. Rougier, N. P., Noelle, D. C., Braver, T. S., Cohen, J. D. & O’Reilly, R. C. Prefrontal cortex and flexible cognitive control: rules without symbols. Proc. Natl. Acad. Sci. U. S. A. 102, 7338–7343 (2005).

51. Flesch, T., Nagy, D. G., Saxe, A. & Summerfield, C. Modelling continual learning in humans with Hebbian context gating and exponentially decaying task signals. PLoS Comput. Biol. 19, e1010808 (2023).

52. Giallanza, T., Campbell, D. & Cohen, J. D. Toward the emergence of intelligent control: Episodic Generalization and Optimization. Open Mind 8, 688–722 (2024).

53. Russin, J., Zolfaghar, M., Park, S. A., Boorman, E. & O’Reilly, R. C. A neural network model of continual learning with cognitive control. CogSci 44, 1064–1071 (2022).

54. Bell, A. M., Hankison, S. J. & Laskowski, K. L. The repeatability of behaviour: a meta-analysis. Anim. Behav. 77, 771–783 (2009).

55. Dingemanse, N. J., Both, C., Drent, P. J., van Oers, K. & van Noordwijk, A. J. Repeatability and heritability of exploratory behaviour in great tits from the wild. Anim. Behav. 64, 929–938 (2002).

56. Mkrtchian, A., Valton, V. & Roiser, J. P. Reliability of decision-making and reinforcement learning computational parameters. Comput. Psychiatr. 7, 30–46 (2023).

57. Sih, A. & Del Giudice, M. Linking behavioural syndromes and cognition: a behavioural ecology perspective. Philos. Trans. R. Soc. Lond. B Biol. Sci. 367, 2762–2772 (2012).

58. Kelly, C. A. & Sharot, T. Individual differences in information-seeking. Nat. Commun. 12, 7062 (2021).

59. Webster, D. M. & Kruglanski, A. W. Individual differences in need for cognitive closure. J. Pers. Soc. Psychol. 67, 1049–1062 (1994).

60. Roets, A. & Van Hiel, A. Separating ability from need: clarifying the dimensional structure of the Need for Closure Scale. Pers. Soc. Psychol. Bull. 33, 266–280 (2007).

61. Condon, D. M. & Revelle, W. The international cognitive ability resource: Development and initial validation of a public-domain measure. Intelligence 43, 52–64 (2014).

62. Dworak, E. M., Revelle, W., Doebler, P. & Condon, D. M. Using the International Cognitive Ability Resource as an open source tool to explore individual differences in cognitive ability. Pers. Individ. Dif. 169, 109906 (2021).

63. Lee, J. K., Rouault, M. & Wyart, V. Compulsivity is linked to suboptimal choice variability but unaltered reinforcement learning under uncertainty. Nat. Ment. Health 3, 229–241 (2025).

64. Acerbi, L. & Ji, W. Practical Bayesian optimization for model fitting with Bayesian adaptive direct search. Neural Inf Process Syst 30, 1836–1846 (2017).

65. Stephan, K. E., Penny, W. D., Daunizeau, J., Moran, R. J. & Friston, K. J. Bayesian model selection for group studies. Neuroimage 46, 1004–1017 (2009).

66. Sutton, R., McAllester, D. A., Singh, S. & Mansour, Y. Policy gradient methods for reinforcement learning with function approximation. Neural Inf Process Syst 12, 1057–1063 (1999).

